# Magnetic molecular nanorulers enable combinatorial biosensing and determine distances at the molecular level

**DOI:** 10.64898/2026.10.19.760221

**Authors:** Yihao Wang, Florian Wolgast, Tim Schröder, Rabia Amin, Caroline Körösy, Meinhard Schilling, Philip Tinnefeld, Jan Lipfert, Thilo Viereck, Aidin Lak

**Affiliations:** Institute for Electrical Measurement Science and Fundamental Electrical Engineering and Laboratory for Emerging Nanometrology (LENA), Hans-Sommer-Str. 66, Braunschweig 38106, Germany; Department of Chemistry and Center for NanoScience, LMU Munich, 81377 Munich, Germany; Soft Condensed Matter and Biophysics, Department of Physics and Debye Institute for Nanomaterials Science, Utrecht University, The Netherlands; Institute of Physics, University of Augsburg, 86159 Augsburg, Germany

**Keywords:** Ultra-responsive magnetic nanoparticles, Engineered magnetization dynamics, Molecular nanorulers, Combinatorial biosensing, Molecular distances, Magnetic particle spectroscopy

## Abstract

Molecular nanorulers are essential tools for mapping intramolecular distances and studying nucleic acid and protein interactions. Here, we present MARs (**MA**gnetic molecular nano**R**ulers), a fundamentally new class of nanorulers based on measuring the magnetization dynamics of rationally engineered and ultra-responsive colloidal magnetic nanoparticles with magnetic particle spectrometry (MPS). By tuning the grafting density of DNA receptors on MARs, we reveal surface molecular crowding as a simple, yet programmable parameter to tune the response of MARs over a broad range of target DNA concentrations. We show that MARs can distinguish target binding positions along receptor strands by regulating how much the binding effect is translated into the response. Combining their switchable binding-to-response and DNA hairpins, we show that MARs can resolve the full combinatorial space of four short DNA targets (2^4^-1) in a one-design-for-all fashion, with blind assays achieving 80% best-prediction accuracy. By modelling MPS response of MARs using the Fokker-Planck equation, we demonstrate that their response is well predictable. Using biotinylated DNA constructs and calibrating their lengths for different numbers of base pair, we reveal that MARs can determine the distance between bound monostreptavidin and their surface with a dynamic distance range of up to 13 nm and a resolution of 1.3 ± 0.2 nm. Our study establishes a new class of molecular nanorulers for resolving conformational changes in nucleic acids and measuring distances at the molecular level in free solution via an adaptable, simple, and rapid workflow, making MARs highly promising for high throughput screening assays.

## Introduction

The biological activities of nucleic acids and proteins are fundamentally defined by their dynamic conformational landscapes, where subtle rearrangements in molecular structure often translate into profound shifts in function. Therefore, measurements of molecular distances and conformational changes are key to dissecting the structure, dynamics, and functions of biological macromolecules.^1–6^ Förster resonance energy transfer (FRET),^7–16^ surface plasmon resonance,^17–19^ graphene energy transfer,^20–22^ and small-angle X-ray scattering^23–25^ have been developed for measuring intermolecular distances at the nanoscale. While in particular FRET has provided invaluable details at the molecular level, it requires labeling with fluorescent markers and sophisticated optical read out.^19,26,27^ Moreover, FRET’s dynamic range is limited to intermolecular distances < 10 nm due to the Förster radius of donor acceptor pairs and the inverse six power law as well as autofluorescence background in complex biological media make data interpretation challenging.^28^

Magnetic nanoparticles and methods offer critical advantages for biological applications.^29–32^ Being responsive to magnetic fields, magnetic nanomaterials serve as theranostic agents for diagnostics and targeted drug delivery.^33–35^ Magnetic nanoparticles (MNPs) enable so-called “mix & measure” bioassays on crude biological samples thanks to minimal magnetic signal attenuation with depth.^36,37^ Among different magnetic characteristics of colloidal MNPs, their Brownian relaxation time *τ*_B_ changes highly specifically in response to environmental cues such as viscosity, temperature, and ligand-receptor binding on particle surfaces.^31,38–42^ Cryogenic superconducting, nuclear magnetic resonance spectroscopy, and magnetorelaxometry have been used to monitor target-receptor binding on MNPs.^43–48^ Recently, magnetic particle spectroscopy (MPS), which is based on the non-linear magnetization response of colloidal MNPs, has become a rapid, robust, and cost-effective tool for reading out magnetization dynamics of MNPs in solution.^35,40,49–52^ To resolve nanometer changes in particle hydrodynamic size (*D*_h_) due to molecular binding by MPS, engineered MNPs with high magnetic moment *m* (proportional to particle core size *D*_c_) and small *D*_h_ are needed (Eq. 8 and 9 in SI).^53^ Due to the cubic dependence of *τ*_B_ on *D*_h_, small *D*_h_ are crucial for measuring small changes in *D*_h_ by MPS.^54^ For instance, a 0.5 nm change in *D*_h_ would correspond to a ≈ 3% relative MPS signal change for MNPs with a *D*_h_ ≈ 50 nm. Yet, as the MPS signal amplitude increases with *m*, enhancing the signal requires increasing *D*_c_, demanding an ultrathin polymeric shell around MNPs to keep *D*_h_ small. The combination of high *m* and small *D*_h_ is not available commercially, as commercial MNPs often have a multi-core structure (i.e. aggregates of nano-crystallites) that is embedded in a fairly thick organic shell.^31,48,55–57^

Changes in MPS harmonics spectrum in response to target-receptor binding on MNPs have so far been analyzed by considering a simple Brownian motion model, in which *τ*_B_ scales linearly with the particle hydrodynamic volume. The model, however, is not able to decouple the effect of target’s molecular weight from its spatial binding position on the MNP magnetization dynamics and MPS harmonics. Identifying their contributions to the MPS signal change is fundamental for advancing toward high-resolution magnetic molecular probes, as recent studies have shown that the physiochemical characteristics of short nucleic acids are modulated by the microenvironment they experience on nanoparticles.^58–62^ The melting temperature, folding, conformation, and end-to-end distance of ssDNA and dsDNA on nanoparticles differ from those in free solution due to molecular crowding, configuration entropy, and steric repulsions.

Here we present a combination of rationally engineered MNPs and frequency-tailored MPS that provides a new class of magnetic molecular nanorulers (MARs). Our MARs are capable of resolving nanometer length changes in nucleic acid conformations in free solution within few minutes and exhibit a tunable dynamic range and response window depending on ssDNA receptor grafting density. In high density MARs, ssDNA receptors form a brush-like conformation with a characteristic length of ≈ 25-29 nucleotides (nt), enabling them to resolve the spatial binding position of target DNA. Using the ability of MRAs in regulating how much the binding effect is reflected in the MPS response, we demonstrate a full combinatorial sensing of four DNA targets in a one-design-for-all fashion. Labeling MARs with biotinylated double-stranded DNA and calibrating its length, we demonstrate that MARs can measure the distance between bound monostreptavidin and their surface with a dynamic distance range of up to ≈13 nm and a resolution of 1.3 ± 0.2 nm. Our innovative MARs enable adaptable, simple, rapid, and low-cost assays for screening conformational changes in nucleic acids and measuring distances at the molecular level in free solution.

## Results and Discussion

### MARs distinguish between coiled and brush-coiled conformations of ssDNA receptors

We construct MARs by coating our custom Co_0.4_Zn_0.2_Fe_2.4_O_4_ MNPs^29^ with a mean core size of 17.9 ± 2.9 nm (n = 138, Fig. S1) with poly(maleic anhydride-alt-1-octadecene)-PEG(3)-azide polymer and fractionating them using agarose gel (experimental details in Supporting Information (SI)).^58^ The particle size (core + polymer shell) increases to 25.5 ± 1.9 nm (n = 85) after polymer coating, corresponding to a polymer shell thickness of ≈ 3.8 nm, as determined from the analysis of negative-stain TEM images (Fig. 1a, 1b). This specific size increase indicates the nature of MARs being single-polymer-encapsulated MNPs.

**Figure 1.**
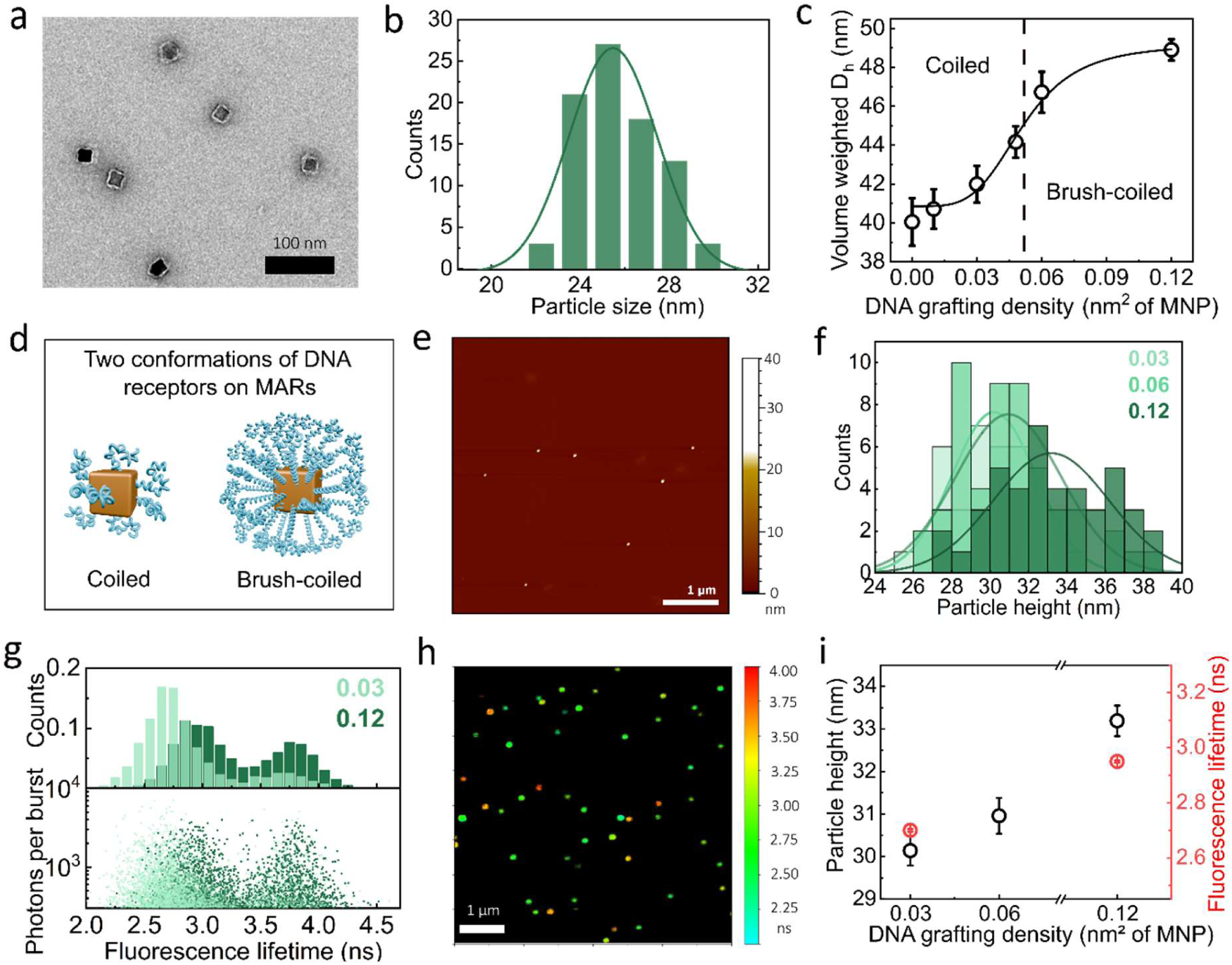
Physicochemical characteristics of MARs at different ssDNA receptor grafting densities. (a) Negative-stain transmission electron microscopy (TEM) image of MARs after poly(maleic anhydride-alt-1-octadecene) (PMAO)-PEG(3)-azide coating and DNA labeling with the receptor 39-1. (b) Particle size histogram of MARs obtained from TEM analysis of the sample in (a). (c) The volume-weighted particle hydrodynamic size (*D*_h_) of MARs labelled with 39 nt ssDNA receptors (Table S2) at different grafting densities, measured by dynamic light scattering (DLS). The dashed line indicates a transition in ssDNA conformation from being coiled to brush-coiled. Data points are the mean ± standard error of the mean (SEM) (n =3) and the solid line indicates a fit with the Hill-like equation (Eq. 3, SI). The DLS measurements were performed at a particle concentration of ≈ 2 nM. (d) Schematic illustrations of MARs with coiled and brush-coiled ssDNA conformation at low and high grafting density, respectively. (e) Atomic force microscopy (AFM) image of MARs at 0.12 ssDNA/nm², drop-cast and imaged in tapping mode. (f) Particle height histograms from AFM imaging (as shown in panel e and Fig. S3) of MARs at different grafting densities. The AFM images (Fig. S2, SI) were acquired in tapping operation mode. (g) Fluorescence lifetime distribution of ATTO643 dyes versus photons per burst from in solution diffusion MARs. The dyes are coupled to MARs via ssDNA labeling at grafting densities of 0.03 and 0.12 ssDNA/nm². The upper histogram shows the distribution of recorded fluorescence lifetimes of individual bursts, normalized to the total number of bursts. (h) Fluorescence Lifetime Imaging Microscopy (FLIM) image of immobilized MARs, coupled to ATTO643 dye-labeled ssDNA at 0.12 ssDNA/nm² density. The color scale represents the fluorescence lifetime of dyes in nanosecond. An upper intensity threshold of 100 photons per pixel was applied. (i) Particle height determined from AFM and fluorescence lifetime versus the DNA grafting density. Data are shown as the mean ± SEM (n = 3).

We then label MARs with 39 nt ssDNA receptor strands (receptor 39-1, Table S2, SI) using the copper-free click conjugation chemistry^58^ at different grafting densities and investigate how the conformation of ssDNA receptors depends on the grafting density. We find that the number of ssDNA per MAR (and thus the actual DNA grafting density) increases linearly with the amount of added DNA in the labeling reaction (i.e. with the nominal DNA grafting density), as determined from depletion assays (Fig. S2, Table S1). However, the particle hydrodynamic diameter *D*_h_ values (as measured by dynamic light scattering) increase non-linearly with the grafting density and follow a Hill-like dependence (Fig. 1c).

At 0.01 ssDNA/nm^2^ grafting density (≈ 20 ssDNA/MAR), *D*_h_ increases by only ≈ 0.7 nm compared to no DNA labeling, suggesting a DNA shell thickness of ≈ 0.35 nm. This increase is within the order of the persistence length of ssDNA which is 0.9-3 nm,^63–65^ suggesting a coiled conformation of ssDNA on MARs at this density. In contrast, *D*_h_ increases by ≈ 8.8 nm (i.e. ≈ 4.4 nm DNA shell) at 0.12 ssDNA/nm^2^ grafting density of (≈ 200 ssDNA/MAR), strongly suggesting the formation of a DNA brush conformation up to a certain length at high density (Fig. 1d).

Next, we examined if these two characteristic DNA conformations can be identified by atomic force microscopy (AFM) imaging (AFM images in Fig. 1e and Fig. S2). The particle height analysis of AFM images of drop-casted MARs shows a size increase from 30.2 to 33.2 nm upon increasing the density from 0.03 to 0.12 (Fig. 1f), which indicates a size increase of 4.7 (i.e. 2.3 nm DNA shell) and 7.7 nm (i.e. 3.8 nm DNA shell) compared to unlabeled MARs (i.e. 25.5 nm), respectively (Fig. 1a). While AFM measures the DNA shell thickness on dried MARs, at 0.12 density, the relative changes in particle size measured by DLS and AFM are comparable, suggesting strongly that ssDNA strands are well stretched out from particle surface at high density and preserve this conformation to large extent after drying. Taken together, DLS and AFM results suggest that at a grafting density > 0.06, ssDNA strands are straightened out due to inter-strand electrostatic repulsions and molecular crowding and form a brush-like shell on MARs.

To further analyze the ssDNA conformations, we characterized MARs at 0.03 and 0.12 densities using single particle fluorescence microscopy and Time Correlated Single Photon Counting (TCSPC) by labeling them with, on average, one ATTO643 fluorescent dye by adding 10% dye-conjugated ssDNA to the labeling reaction (Fig. 1g-i). In solution experiments, the single burst analysis of MARs particles reveals two lifetime populations for both densities (Fig. 1g) which becomes apparent in Fluorescence Lifetime Imaging Microscopy (FLIM) of immobilized MARs particles as depicted in Fig. 1h. The green fluorescence spots resemble the quenched fluorescence lifetime particles, whereas the orange particles show the unquenched fluorescence lifetime of 3.7 ns. The short-lifetime population indicates a distance-dependent quenching effect of the MARs’ surface, with the mean fluorescence lifetime shifting from 2.70 to 2.95 ns as the DNA density increases (Fig. 1i, S4, S5). This is a strong evidence of shorter end-to-end distance of ssDNA strands on MARs at low density than high density. Moreover, we observed a shift in the population distribution: The quenched population dominates at low density, while the unquenched population increases significantly at high grafting density, further supporting our conclusion. To further investigate the origin of two-population distribution, fluorescence correlation spectroscopy (FCS) was employed, which reveals distinct hydrodynamic profiles of both populations (Fig. S5): The quenched population exhibits a larger *D*_h_ (≈ 56 nm) compared to the unquenched population (*D*_h_ ≈ 36 nm, Eq.1 in SI). Therefore, we attribute the unquenched lifetime population to dye-labeled polymeric micelles, possible byproducts of polymer coating. Taken together, our results demonstrate that by increasing DNA grafting density its conformation transits from coiled to brush-coiled.

### Broad binding response window at low receptor density versus sharp binding response at high receptor density

Next, we aimed to understand whether the ssDNA receptor conformation has implications for sensing target DNA in free solution. To tackle this question, we designed hybridization assays wherein fully complementary target DNA is added to ssDNA-MARs (Fig. 2a). We then performed titration assays by measuring MPS spectra as a function of target concentration for MARs with different grafting densities (target 39-1, Table S1). From the MPS harmonics spectra, we computed and analyzed the MPS harmonic ratio H5^th^/H3^rd^ (HR53), a particle concentration-independent metric (see SI for details),^51^ where a drop in the HR53 signal corresponds to an increase in *D*_h_.^40^ The resulting titration curves (Fig. 2a-e) are well-described by a Hill-like model and from the fits we extract two characteristic parameters: (1) the apparent affinity *K*_d_, representing the target concentration at which half of the maximal signal change is achieved and (2) the Hill coefficient *n*_H_, characterizing the cooperativity of the response (Fig. 2f). By increasing the receptor density from 0.01 to 0.12, the apparent *K*_d_ increases from 2.48 to 57.2 nM, indicating a higher binding affinity at lower surface density. We attribute this mainly to the nature of nanoparticle-based affinity assays, where overcoming the steric, electrostatic, and osmotic barriers at high brush density reduces the binding accessibility and efficiency. As a result, the apparent *K*_d_ values obtained from our hybridization assays for a 34 nt target are significantly higher than the intrinsic *K_d_* (≈ 1.4×10^-27^ nM at 25°C, determined from NUPACK simulations) of this specific DNA length in free solution across all DNA densities tested here.^66,67^

**Figure 2.**
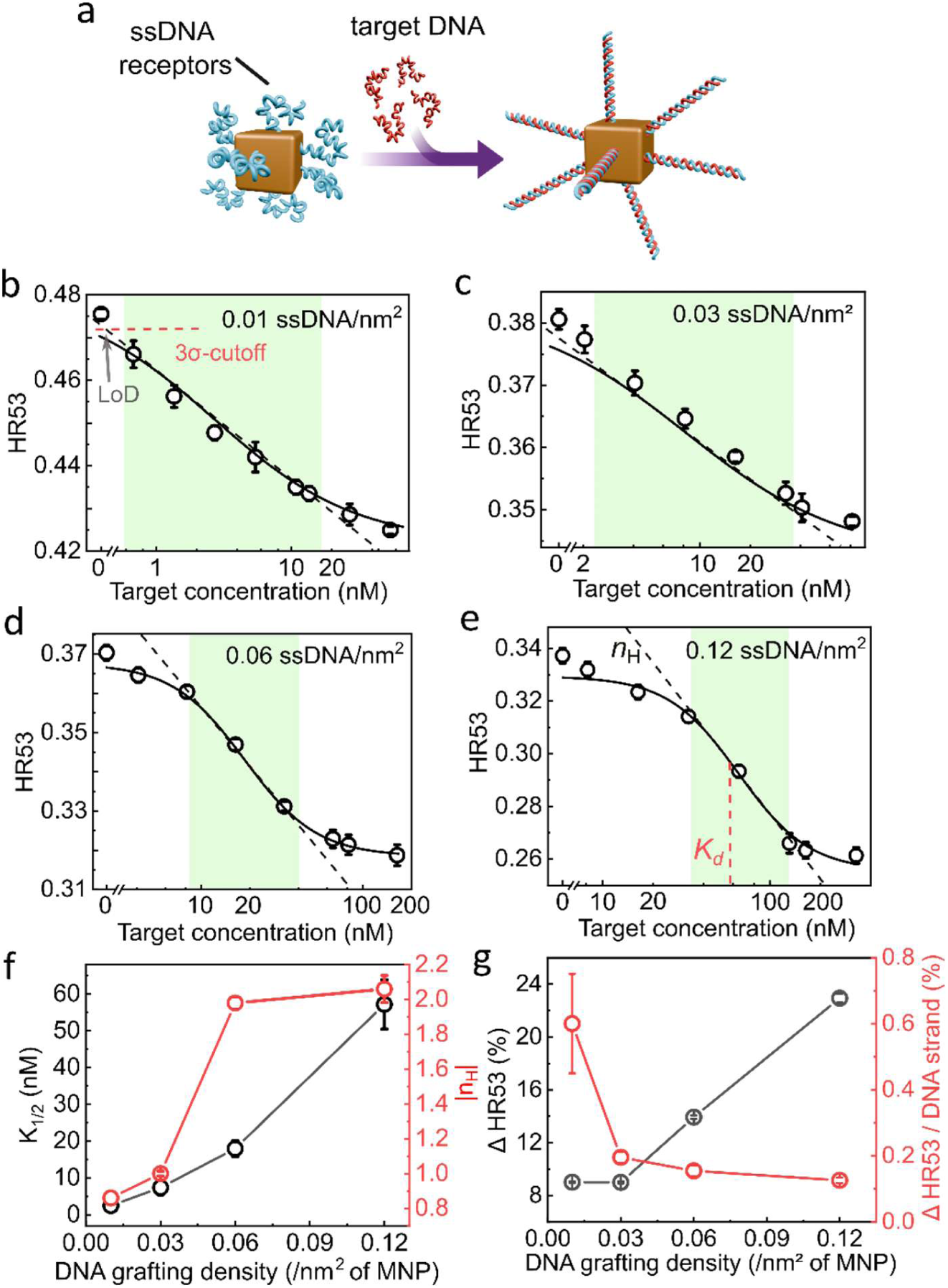
Tuning the MPS response window of MARs by changing ssDNA receptor grafting density and conformation. (a) Scheme of the DNA hybridization assay. Target DNA in solution binds to the receptor DNA on MARs, changing their hydrodynamic radius. (b)-(e) HR53 signal as a function of target DNA concentration for MARs with different ssDNA receptor grafting densities (receptor 39-1, Table S2). The HR53 is the ratio of fifth to third MPS harmonics and is computed from the MPS spectrum. Black lines are fits of the Hill-like equation (Eq. 4, SI). Shaded green areas highlight the response windows and are determined from the overlap between the best linear fit (dashed lines) and the data. The red dashed line in (b) shows the 3σ-cutoff line of the control sample (without target DNA), which is used to determine the limit of detection (LoD) from its intercept with the best linear fit. (f) Evolution of *n*_H_ and *K*_d_ parameters versus the grafting density. (g) The relative change ΔHR53 (Eq. 5, SI) and its normalization to the number of grafted DNA strand (Table S1) versus the grafting density. Data are the mean values of three independent biological replicates, with error bars corresponding to SEM. The samples have a typical particle concentration of 1.5 nM. The excitation field amplitude and frequency of MPS are 15 mT and 1954 Hz.

Compared to the intrinsic *K_d_*, the relatively high apparent *K*_d_ at low grafting density can be partially attributed to the configurational entropy penalty, wherein binding one end of ssDNA receptors to MARs restricts their motional freedom as compared with those in free solution. The brush-like conformation and crowding of ssDNA receptors at high density introduce more configurational and osmotic entropy penalties, which suppress the hybridization and account for much larger apparent *K*_d_ than that of low density. Unlike free solution, where DNA strands are sparsely distributed throughout the entire volume, on the particle surface, only limited effective interaction volumes are available for hybridization, thus further limiting the binding accessibility.^17,68–70^ Looking at the MPS signal gain per receptor, we can support our outlined discussion further (Fig. 2g). The signal gain per receptor decreases with the grafting density, clearly indicating a better binding accessibility and affinity toward target DNA at low than high density.^68^ Considering that the nominal footprint of a DNA duplex on surface is ≈ 13 nm²/molecule^70^ and the interaction volume between a receptor and a target DNA at 0.12 density is 8.7 nm², there is a strong steric and electrostatic hinderance against hybridization, further accounting for the increasing apparent *K*_d_ with density. These phenomena, collectively, lead to higher *K_d_* values in nanoparticle-based affinity assays than bulk methods.^68–71^

The Hill coefficient *n*_H_ values show a similar trend. By increasing the receptor density, *n*_H_ increases from 0.86 to 2.0, showing a clear transition from a non-cooperative to a highly cooperative system. We attribute the cooperativity to molecular crowding, brush-like conformation, and electrostatic effects between neighboring ssDNA receptors,^5,60,72^ resulting in a sharp response (Fig. 2d, 2e). Whereas at low density ssDNA receptors experience weak electrostatic effect, strong repulsive interactions are present in highly crowded DNA brush at high density, leading to a cooperative response.^5^ We find that grafting density strongly modulates assay sensitivity: MARs at low grafting density are sensitive to DNA concentrations < 1 nM by undergoing a coiled-to-duplex conformational change (Fig. 2b). In contrast, at high grafting density, MARs respond sharply to 10-100 nM target concentration, where cooperativity governs the sensitivity (Fig. 2d, 2e). Taken together, our results clearly demonstrate that the DNA hybridization on MARs is strongly modulated by organization of ssDNA receptors on the nanoscale on particle surface. Additionally, we observed that the MARs dynamic response window exhibits two distinct behaviors depending on the grafting density. At 0.01 and 0.03 density, the titration curves show a response window spanning from 0.5 to 16 nM and 2.7 to 37 nM, respectively, which corresponds to nearly 1.5 orders of magnitude dynamic range (shaded areas in Fig. 2b, 2c). At 0.01 density, we determined a limit of detection (LoD) of 315 pM. At 0.06 and 0.12 density, the dynamic response window spans from 9 to 42 nM and 32 to 128 nM, revealing less than one order of magnitude dynamic range and a much sharper response (Fig. 2d, 2e). Homogeneous magnetic bioassays have typically one order of magnitude dose-response dynamic range.^73^ In earlier works, a broad dynamic range was realized only by combining measurements at different MNP concentrations.^74^ Our results establish receptor conformation and binding accessibility as programmable parameters to expand the response range and demonstrate up to 1.5 orders of magnitude dynamical range in a one-pot assay.

### MARs can distinguish the spatial binding position of target DNA on receptors

We then wondered if we can distinguish the spatial binding position of target DNA along the receptor in our MAR assay (Fig. 3a). Our hypothesis is that the DNA brush should have a characteristic brush length (*L*_B_), with base pairs further away from *L*_B_ coiling randomly, that plays a role here. We propose that depending on whether target DNA binds to the brush or coiled segment of the receptor, the binding is coupled to MARs magnetization dynamics (i.e. HR53 ratio) to different extent.

**Figure 3.**
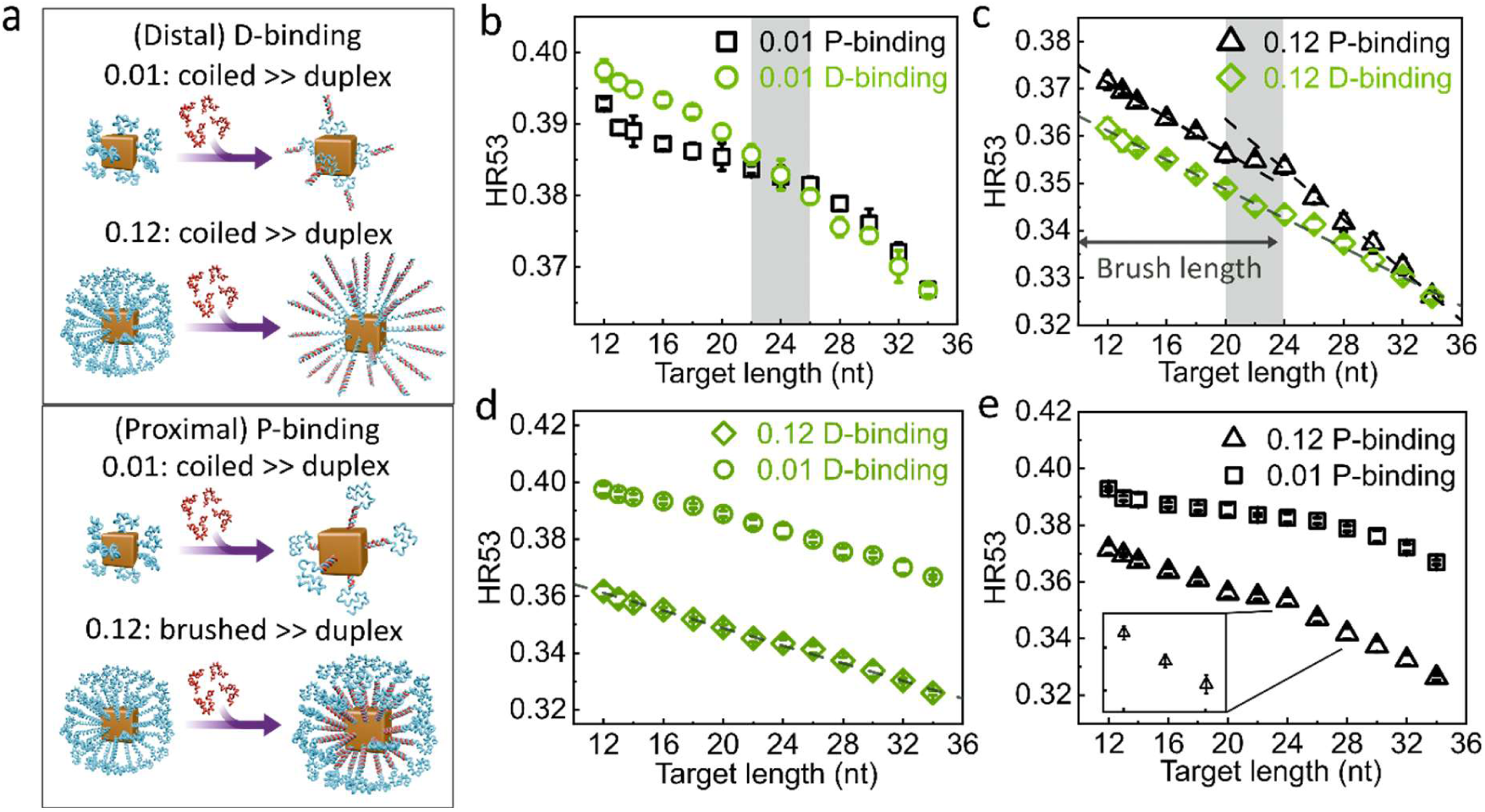
Distinguishing the spatial binding position of target DNA with MARs. (a) Schematic illustrations of target-receptor binding at the distal (D) or proximal (P) end of the receptors on MARs at 0.01 and 0.12 grafting densities. (b)-(e) HR53 changes over the target DNA length up to 34 nt (full hybridization) for the P-and D-binding for both densities. Dashed lines are the best linear fits to data. For the P-binding at 0.12 density, panel (c), two lines with different slopes were used to reconstruct the data over the whole range. The gray shaded area in (b) highlights the convergence of the D-binding and P-binding at ≈ 22-26 nt in case of 0.01 density. The gray marked region in (c) shows the divergence of two linear fits for the P-binding at 0.12 density, highlighting where the coiled to brush ssDNA conformation transition occurs. The inset in panel (e) represents a zoomed-in view of target length 24-28 nt, indicating a 2 base pairs resolution of MARs. Data are shown as the mean ± SEM (n=3) with some error bars smaller than symbols. The measurements were performed at a typical particle concentration of 1.5 nM. Each data set was obtained on the same DNA-labelled MARs, indicating the same number of DNA receptors per MAR.

To test our hypothesis, we labelled MARs with 34 nt ssDNA receptors at 0.01 and 0.12 densities (receptor 39-2, having 5 nt spacer, a slightly modified version of the receptor 39-1 to avoid secondary structures,^67^ Table S2) and varied the target DNA length from 12 to 34 nt, in steps of two base pairs. We designed two sets of experiments: (1) target binding to the distal end (D-binding, i.e. to the end away from the surface) or (2) to the proximal end (P-binding, i.e. to the end closest to the particle surface) of the receptor (Fig. 3a). For all the conditions, regardless of the density and the binding position, HR53 decreases with increasing target length (Figure 3b-d), indicating that a transition from ssDNA to dsDNA retards the MARs magnetization dynamics. For both densities, the HR53 values for P- and D-binding fall on top of each other for the target of 34 nt length (Fig. 3b, c), as expected, since for the fully complementary strand there is no difference between P- and D-binding, and highlighting the repeatability of our assays (see Fig. S6 for DLS results).

Interestingly, for shorter target DNA strands the grafting density plays a critical role. At low grafting density, P-binding exhibits lower HR53 values than D-binding for targets < 22-26 nt (Fig. 3b). According to our earlier discussion, this is where the DNA receptors are coiled and sparsely populated on MARs (Fig. 3a). We argue that, in this configuration, the binding to the proximal end of the receptor is coupled more efficiently to the particle’s dynamics than the binding to the distal end, leading to a larger drop in HR53 (Fig. 3b). We attribute this to the presence of a flanking ssDNA segment ≥ 12-16 nt between the binding position and particle surface that weakens the transmission of the DNA binding effect to particle dynamics (HR53) (scheme in Fig. 3a and Fig 3b). The HR53 trend changes completely for target lengths > 24 nt. In this regime, the HR53 values are essentially indistinguishable within measurement uncertainties (Fig. 3b). According to our proposed concept and in case of D-binding, this is where the ssDNA segment between the dsDNA segment (i.e. where target is bound) and the particle core is short enough to couple the binding-to-dynamics efficiently. Our results show that low receptor density is favorable for resolving the binding position of DNA target using MARs. For both P- and D-binding, the change in the particle molecular weight due to binding is identical for the same target. This means that the signal change is governed by how binding is coupled to dynamics rather than changes to the particle molecular weight.

At high density, the behavior of MARs in response to where the target is bound is very different. In contrast to low density, D-binding shows lower HR53 values than P-binding throughout the whole range, converging only for the full 34 nt target DNA. In case of P-binding, the HR53 values seem not to change much between 20 and 24 nt, after which they drop with a higher rate than before (see dashed fit lines), as if duplexing beyond this range leads to more changes in the relaxation dynamics (Fig. 3c, shaded area). We argue that this shift stems from a brush to coiled transition at ≈ 20-24 nt range (corresponding to 25-29 nt considering 5 nt adenine spacer, Table S2), a so-called characteristic brush length *L*_B_, up to which ssDNA strands are brushed on MARs.

The fact that P-binding causes less drop in the HR53 values than D-binding indicates the critical role of the DNA brush and *L*_B_ in signal change. In case of P-binding and for targets ≤ *L*_B_, as the receptor strands are brushed, duplexing causes a so-called brushed-to-duplex conformational change. In case of D-binding, the situation is quite different. For targets ≥ (34 - *L*_B_), a so-called coiled-to-duplex conformational change occurs, a more drastic change than the brushed-to-duplex, slowing down the Brownian relaxation and dropping HR53 more. This effect is also reflected in a more rapid drop of HR53 in P-binding compared to D-binding for targets ≥ *L*_B_, where the coiled-to-duplex transition governs (Fig. 3c). Over the whole target length range and in the P-binding, the binding is efficiently coupled to the MARs’ dynamics through the brush and in the absence of any flanking segment. Thanks to the efficient transmission of binding to dynamics, MARs exhibit a 2 bp spatial resolution at P-binding for targets > *L*_B_ (Fig. 3e inset). Regardless of the target binding position, high density MARs exhibit a larger drop in HR53 upon target binding than low density MARs (Fig. 3d, 3e), which can be attributed to more ssDNA per MAR, giving rise to more DNA hybridization and overall to a larger effect on the HR53 signal.

### MARs enable dual-plex sensing by harnessing coupled-decoupled binding-to-dynamics concept

Next, we were interested to know if we can develop multiplexed assays by exploiting the unique differences in how DNA binding is transmitted to particle dynamics depending on its binding location. First, we designed a linear DNA receptor with two 15 nt recognition domains, one at the proximal (P) and one at the distal (D) end with respect to MRAs surface (2D-receptor, Table S3, Fig. 4a). We then added targets in different combinations: one target complementary only to domain P (P*) or to domain D (D*) or both targets and measured MPS spectrum in a mix & measure fashion (Fig. 4b). As expected, upon P* binding to P, the HR53 drops compared to the control sample (Fig. 4c, 0T). The signal drop is larger in case of D* than for P*, indicating that distal binding has a larger effect on particles dynamics following our discussed coiled-to-duplex concept. Upon adding both targets to the test tube, HR53 drops further since the total length of target DNA bound is now 30 nt (Fig. 4c, 0T). Taken together, these data clearly show that we can systematically distinguish P*, D* and the presence of both targets in a one-pot assay with high statistical significance (p < 0.001, Fig. 4c, 0T).

**Figure 4.**
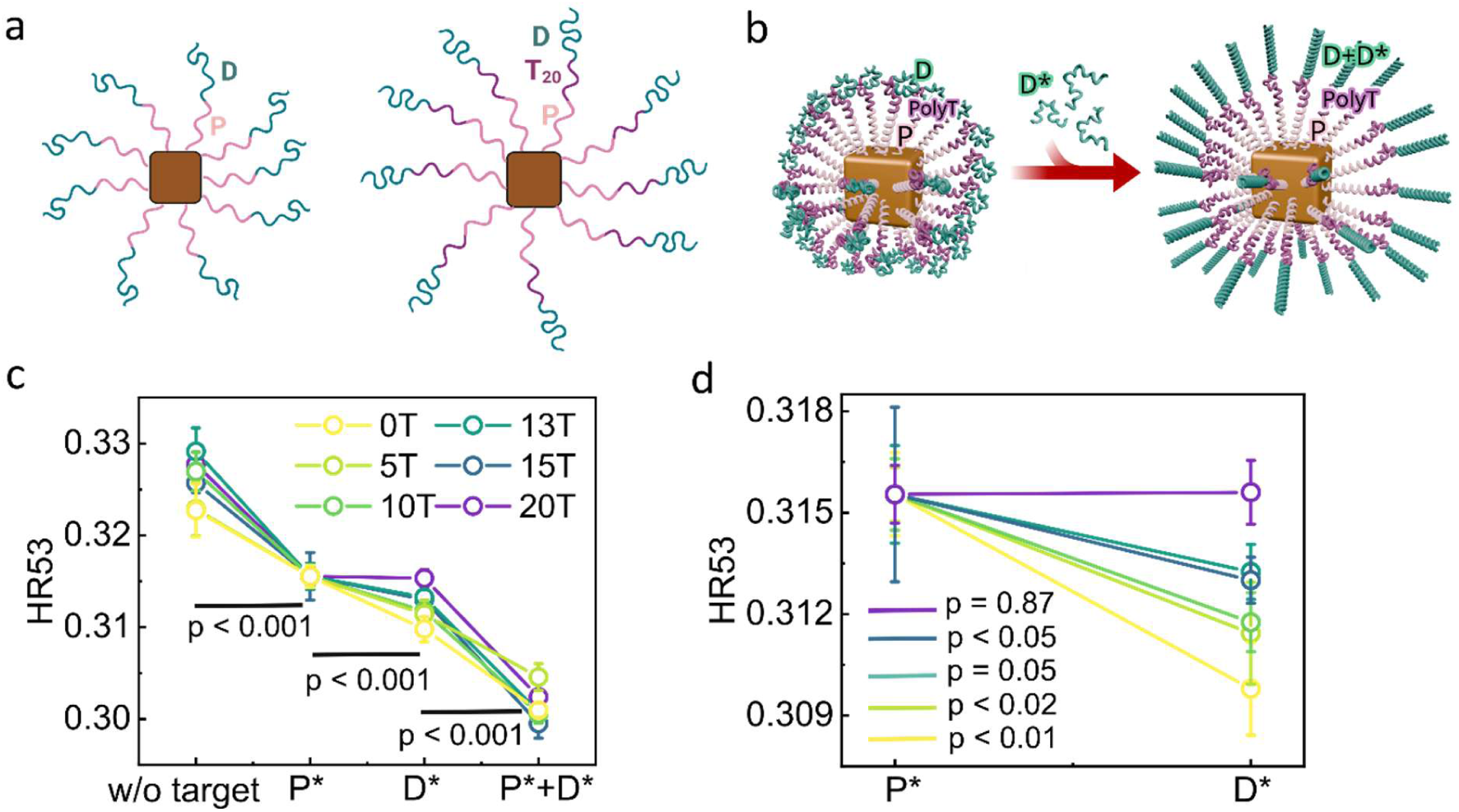
Dual-plex sensing of short DNA sequences by harnessing coupled-decoupled binding-to-dynamics. (a) Schematic representation of high-density MARs (0.12 ssDNA/nm²) with 2D-receptor and modified 2D-receptors by adding dT segment between P and D domains (dT_20_ is shown exemplarily). (b) 3D illustration of MARs at 0.12 ssDNA/nm² density with 2D-receptor DNA upon sensing target D*. The purple segment indicates the poly(T) between two domains. (c) HR53 signal of MARs with 2D-receptor with different poly(T) lengths and for different target sets. Data of different poly(T) lengths is normalized to D* of 0T for better comparison. P-values were determined between adjacent target variations shown at x axis based on the unpaired two-tailed t-test. (d) Zoomed-in view of P* and D* target binding responses from (c). Data points are the mean ± SEM (n = 6 independent biological replicates). The samples have a typical particle concentration of 2 nM. The p-values were determined based on the unpaired two-tailed t-test.

Next, we explored if having a flanking ssDNA segment between the two recognition domains changes that way these assays work, building on our previous findings showing that the binding-to-dynamics is partially decoupled in the presence of a flexible linker (Fig. 3b). To design such assays, we modified the 2D-receptor by adding a poly-deoxythymidine (dT) segment of varying lengths (5-20 nt) between P and D domains (Fig. 4a-b). As the poly(T) linker length increases, the effect of D* to D binding on HR53 is less and less distinguishable from P* to P binding (Fig. 4c-d). At the linker length of 20 nt, the HR53 values of both D* or P* binding are essentially identical (Fig. 4d), meaning that MARs are not able to distinguish which target is present. These results clearly indicate a progressive decoupling of binding-to-dynamics by inducing a ∼ > 13 nt flanking ssDNA linker between the recognition coiled domain and the brush domain. According to our results, an efficient binding-to-dynamics coupling is required to have one-pot dual-plex assays using MARs, a feature MARs offer when having no flanking segment.

### Full combinatorial biosensing by combining hairpins and coupled-decoupled dynamics

We went further and aimed to achieve more sophisticated multiplexed assays by leveraging hairpins as recognition domains in combination with the outlined coupled-decoupled dynamics in one-design-for-all fashion. As hairpins undergo a major conformational change from a loop structure to a rigid dsDNA upon binding to their complementary sequence, we reasoned that a hairpin design could provide a larger change in MPS signal than our linear design.^75^ We designed a receptor complex consisting of four 20 nt loop domains (loop-receptor) (Fig. 5a, Fig. S7, Table S3) and performed assays across the full combinatorial space (n = 2^4^ -1) of four targets using MARs with this receptor at 0.12 ssDNA/nm² density. Looking at the combinations with only one target present (Fig. 5b), we observed that binding of L2* to L2 leads to the highest HR53 signal drop among all the single-target combinations. This behavior matches what we observed in the 2D-receptor, indicating again that the most effective binding-to-dynamics coupling occurs when the target binds to a loop that is placed right after the brush domain. It is remarkable to see that by binding a target further away from the brush, the HR53 signal drops less and less systematically (see L3* and L4* targets, Fig. 5b). Consistent with our results on the 2D-receptor with 20 nt spacer, here, there is no significant difference in HR53 between L1* and L3* (Fig. 5d, ns), suggesting the flanking role of L2 loop, hence partially compromising the binding-to-dynamics of L3*.

**Figure 5.**
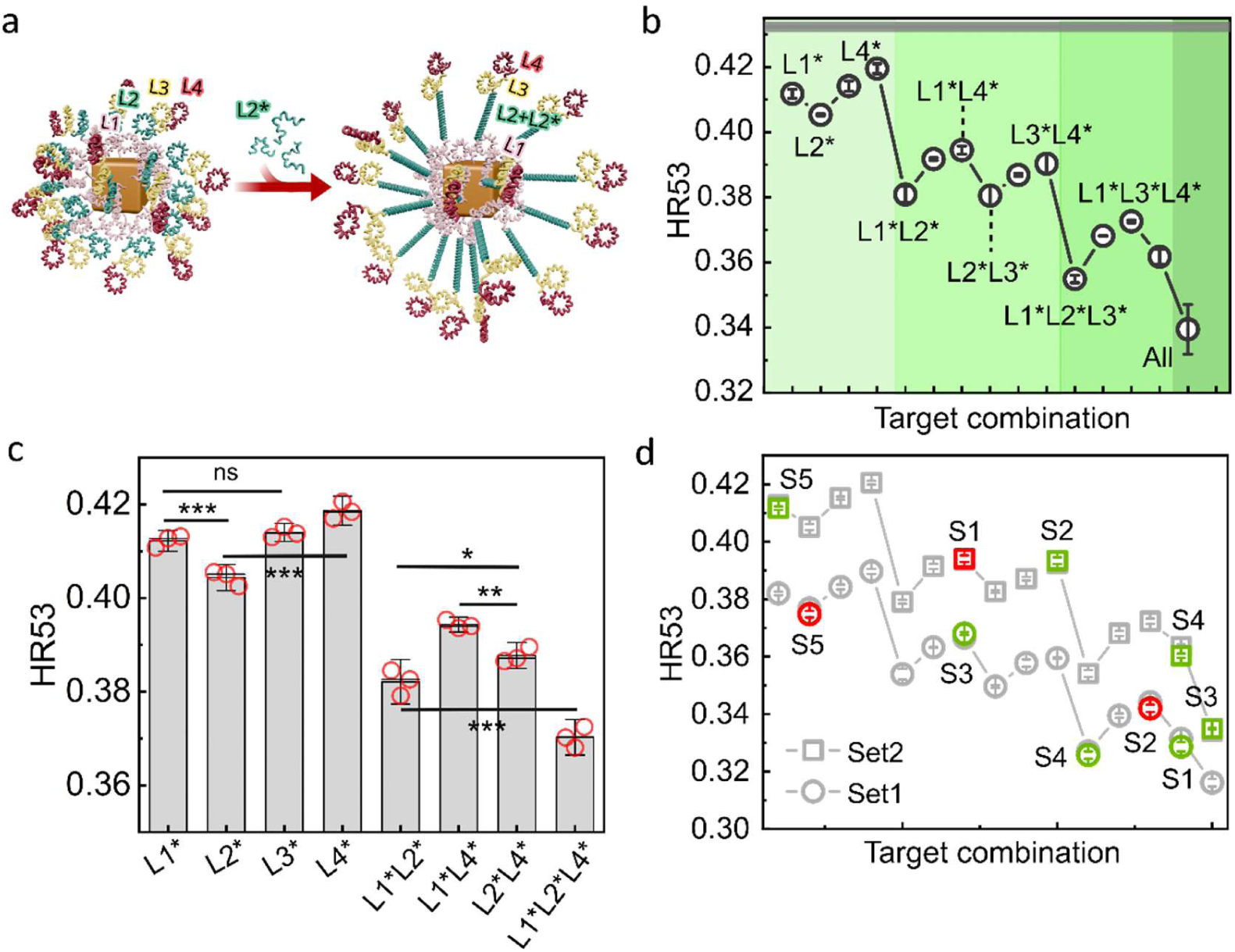
Full combinatorial sensing of four DNA targets by combining hairpins and coupled-decoupled concept. (a) Scheme of high-density MARs (0.12 ssDNA/nm²) with DNA loop-receptors before and after sensing L2* target. (b) HR53 signal of MARs with loop-receptors versus different target combinations. Data represent the means from two independent DNA functionalization sets (set 1, normalized to the control sample without target DNA of set 2 to remove the set-to-set variation, and set 2) for each target combination, with the error bars corresponding to SEM (n=6). The horizontal gray area represents the signal of control sample in the absence of target DNA. The ratio of target DNA to the receptor DNA is 2:1 to ensure binding saturation. Target combinations were categorized into four groups: single, double, triple, and quadruple target sets and highlighted by color gradients from light to dark. Data within each group are labeled in a sequence from L1* to L4*. (c) Zoomed-in view of some exemplarily combinations of L1*, L2* and L4* plus single L3* target binding response from set 2 (d), demonstrating that three targets L1*, L2*, L4* and their specific combinations are with high significance distinguishable (data for set 1 is shown in Fig. S8). The p-values are determined using unpaired two-tailed t-tests (*p < 0.05, **p < 0.01, ***p < 0.001; ns, not significant). Error bars here demonstrate 3SEM (n=3). (d) HR53 signals of whole samples for set1 and set2 and five blind samples mapped onto the respective calibration set. Green data points show the correct predictions and red symbols indicate the wrong predictions. Data in (d) are shown as the mean ± SEM (n = 3). The measurements were performed on samples with a typical particle concentration of 1.5 nM.

Trends we observed for double- and triplet-target combinations are similar to that of the single-target combinations. The largest signal reduction (i.e. largest *D*_h_) appears when the L2* target is present and the lowest signal drop is observed for L4*. These trends can also be understood similarly to the single-target combinations. The highest signal drop appears when the target binds right after the brush domain, while the lowest signal drop is due to the flanking ssDNA segment between the target recognition and the brushed segment. As expected, when all four targets present, the highest signal drop among all the combinations is achieved. Our results clearly demonstrate that MARs are able to determine the number of targets present in unknown samples and distinguish three different targets L1* (binding to the brush), L2* (binding to the segment right after the brush), L4* (binding to the site having a long flanking segment) and their full combinatorial sets with high significance (Fig. 5d).

### MARs can detect four targets in blind combinatorial assays with high accuracy

To move toward testing samples with unknown combinations of targets, we conducted blind tests, wherein five target combinations were chosen randomly from the full combinatorial pool and pipetted into five test tubes (Table 1). After performing two sets of independent blind experiments using our MPS assays, we compared the HR53 values of the blind samples with their corresponding calibration curve that we obtained on known samples (Fig. 5c, Set1 and Set2). Comparing our estimations with the actual target combinations (Table 1), we show that MARs with four loop-receptors can detect four targets with 70% average accuracy in a one-pot and one-design-for-all assay. As discussed above, the L3* target is hard to distinguish from the L1* target, thus lowering the overall accuracy of the blind tests.

**Table 1.** Comparison of estimated versus actual target combination identity in five blind samples.

|  | Set 1 |  | Set 2 |  |
| --- | --- | --- | --- | --- |
|  | Estimated identity | Actual identity | Estimated identity | Actual identity |
| Sample 1 | L2*+L3*+L4* | L2*+L3*+L4* | L1*+L4* | L1*+L3* |
| Sample 2 | L1*+L3*+L4* | L2*+L3* | L3*+L4* | L3*+L4* |
| Sample 3 | L1*+L4* | L1*+L4* | All | All |
| Sample 4 | L1*+L2*+L3* | L1*+L2*+L3* | L2*+L3*+L4* | L2*+L3*+L4* |
| Sample 5 | L2* | L1* | L1* | L1* |

### MARs can determine distances up to 13 nm with nanometer resolution: Experiments and Fokker-Planck simulations

Moving toward using MARs for measuring distances at the molecular level, we aimed to understand how the MAR’s MPS signal varies with the length of grafted molecules. For these assays, we opted to use dsDNA for its rigidity up to its persistence length (≈ 50 nm), making it one of the most studied molecular rulers.^23^ Having labelled MARs with dsDNA with varying numbers of base pairs (bp) at 0.12 ssDNA/nm² density via ssDNA conjugation and subsequent hybridization, we first determined the effective dsDNA shell thickness using DLS. This was done by subtracting the radius of MARs from that of dsDNA-labelled MARs, hereafter called the end-to-end distance of dsDNA (*R*_ee_, Fig. 6). By computing the DNA contour lengths (i.e. *L*_0_ = number of bp × 0.34 nm/bp) for each length and comparing them with the *R*_ee_ values, we observed interesting features. We found that the *R*_ee_ values increase at ≈ 0.34 nm per base pair up to 15 bp, matching perfectly a growth rate expected for *L*_0_ per base pair.^23,76^ This behavior strongly suggests that dsDNA molecules are straightened out perfectly on MARs with no bending up to this length.

**Figure 6.**
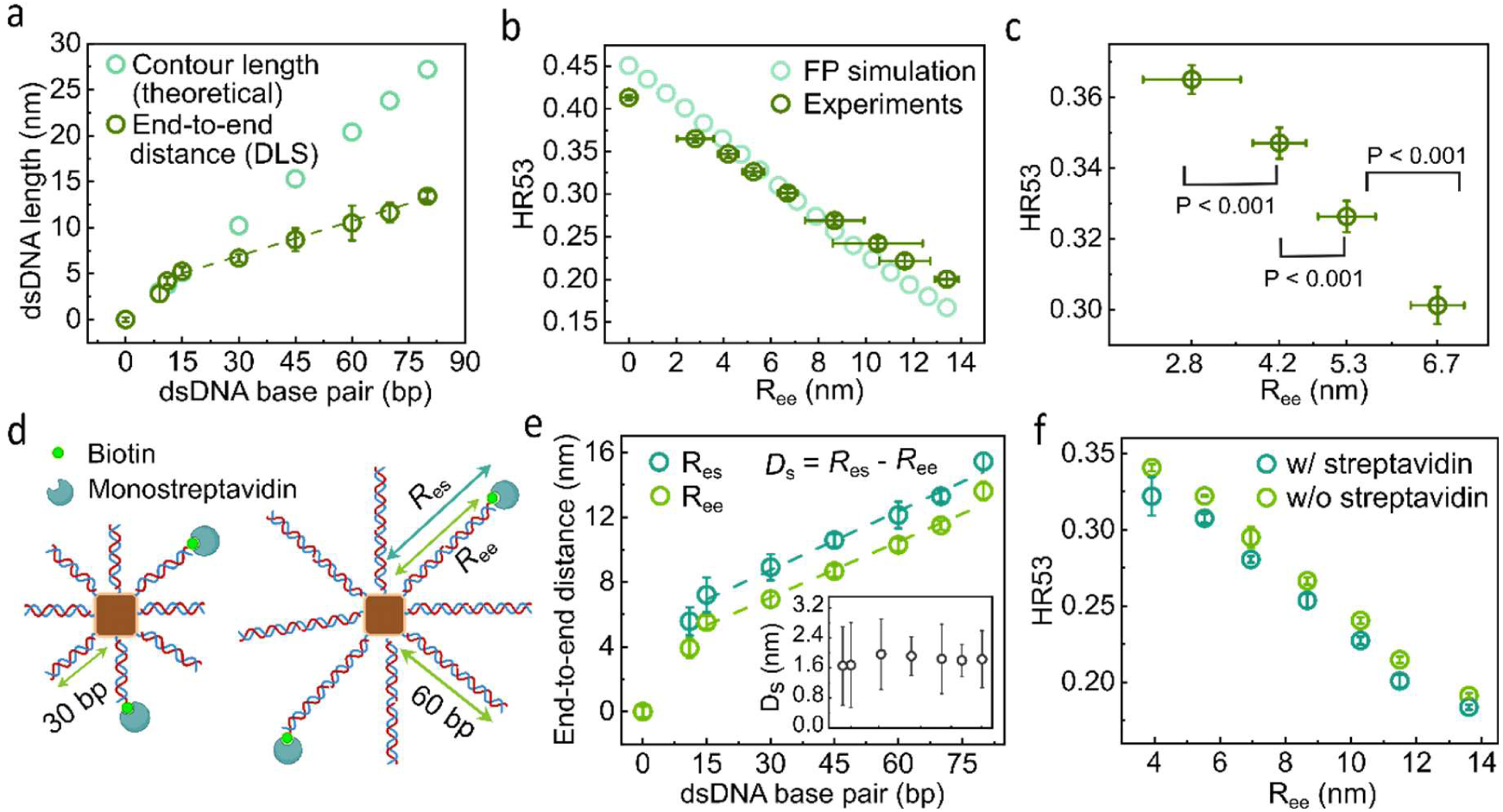
Double-stranded DNA-labelled MARs as molecular nanorulers with > 10 nm dynamic range. (a) Theoretical contour lengths and end-to-end distances (from DLS) of dsDNA on MARs as a function of number of dsDNA base pairs. The dashed line represents the linear fit to data from 15 bp to 80 bp with a slope of 0.13 (R² = 0.99). (b) Experimental (dark green) and using the Fokker-Planck equation (Eq. 6 in SI) computed (light green) HR53 of MARs labelled with dsDNA versus dsDNA end-to-end distance *R*_ee_. (c) Zoomed-in view of HR53 vs. *R*_ee_ for the range from 2.8 to 6.7 nm. The p values were calculated based on unpaired two-tailed t-tests. Data in (b) and (c) are shown as the mean ± SEM (n = 3 independent replicates of DNA labeling). (d) Schematic illustration of molecular distance determination using biotin-streptavidin binding pair as a model system, showing 30 and 60 bp dsDNA as examples with 20% biotin occupancy. *R*_es_ represents the end-to-end distance between MAR surface and the mono-streptavidin molecule. (e) End-to-end distance values before and after binding mono-streptavidin molecules determined from DLS as a function of number of dsDNA base pairs. Both dashed lines represent linear fits to data from 15 bp to 80 bp, yielding slopes of 0.12 (dark green, R² = 0.98) and 0.11 (light green, R² = 0.99), respectively. The inset is the difference between *R*_ee_ and *R*_es_ from 11 to 80 bp, standing for the apparent diameter of mono-streptavidin (*D*_s_). (f) Experimental HR53 of MARs before and after binding mono-streptavidin as a function of *R*_ee_. All measurements were performed on samples at ≈ 1.8 nM particle concentration.

For longer dsDNA constructs, the trend looks different, yet linear. The *R*_ee_ values increase linearly at ≈ 0.13 nm/bp, thus diverging from the theoretical 0.34 nm/bp for *L*_0_ (Fig. 6a).^23^ We attribute the divergence mainly to thermal bending and curving fluctuations governed by worm-like chain (WLC) behavior of dsDNA.^77,78^ As a result, the rather bent and long dsDNA strands reduce the radial extension of the DNA layer, accounting for diverging *R*_ee_ values from the corresponding *L*_0_. Additionally, the DLS hydrodynamic size of MARs is smaller than a simple addition of the theoretical DNA shell thickness to the MAR physical size (i.e. core + polymer shell) due to porous nature of DNA brush, contributing to decreasing the average *R*_ee_ at longer dsDNA. Note that our assays do not enable us to exclude batch-to-batch variability s (≈ 10%) in terms of ssDNA per MAR, as it may well occur for long ssDNA strands due to steric hindrance and electrostatic repulsions.^79,80^

Building upon these new insights, we then measured HR53 values as a function of *R*_ee_ and asked if MARs are able to determine distances at the molecular level by measuring MPS, taking only a minute. We observed a monotonic decrease of HR53 values with *R*_ee_ over a broad range up to ≈ 13 nm, indicating a predictable and computable magnetic response of MARs (Fig. 6b). By doing statistical analyses on these results, we found that they are able to reliably distinguish all adjacent data points (p < 0.01 or 0.001) with the best spatial resolution of 1.3 ± 0.2 nm (Fig. 6c).

Finally, we computed the HR53 values at different dsDNA lengths using the Fokker-Planck (FP) equation, a model describing dynamics of MNPs in solution by solving the time-dependent probability distribution function of magnetic moments in response to external magnetic fields and thermal fluctuations._81_ We performed series of FP simulations by taking the above discussed physicochemical properties of MARs (Fig. 1) and considering *R*_ee_ as the only free parameter (Eq. 9 in SI). The computed HR53 values match closely the experimental values nearly over the whole *R*_ee_ range (Fig. 6b). Small discrepancies between experimental and computed HR53 at distances < 2 nm and >12 nm can be seen which could be attributed to inaccuracies in determining particle’s magnetic moment and simplified assumptions used in the model: (1) mathematical conversion of cubic to spherical particle geometry of same volume, and (3) non-interacting MARs. Nevertheless, both our experimental and computational findings clearly demonstrate the capability of MARs to resolve up to ≈ 13 nm distances from their surface (Fig 6b, 6c), a dynamic range beyond what FRET can achieve (i.e. ≈ 10 nm)^27^.

### MARs can measure the average distance between proteins and their surface: mono-streptavidin-biotin model system

In the following step, we wondered if MRAs can measure the average distance between proteins and their surface. Here and for the proof-of-principle, we studied biotin-mono-streptavidin ligand-receptor pair for its high affinity and binding strength by having biotin groups at the distal end of dsDNA (Fig. 6d). Looking at the DLS results, we found a linear correlation between the end-to-end distance and number of dsDNA base pairs for both biotin (*R*ee) and biotin-streptavidin (*R*_es_) with slopes similar to what we found for dsDNA-MARs (Fig. 6e). These results indicate two things: (1) number of biotin-mono-streptavidin pairs per MAR is for the dsDNA lengths, and (2) binding of streptavidin to MARs changes their translational dynamics linearly over the range studied. To estimate the apparent size of streptavidin, we computed *D*_s_ = *R*_es_ *-R*_ee_ for each DNA length and noted that their mean value is at 1.8 ± 0.1 nm, varying for only ≈ 5% over the whole range (inset of Fig. 6e). Moving on, we looked at how HR53 changes by streptavidin conjugation. We observed a systematic and reproducible drop in HR53 upon biotin-streptavidin pairing over the whole DNA length explored here.

## Conclusions

In this study, we present a fundamentally new class of molecular nanorulers by reading the magnetic response of rationally engineered colloidal MNPs to alternating magnetic fields using a frequency-tailored magnetic particle spectroscopy in free solution. Having a tunable and thin polymer-DNA receptor shell, MARs convert nanometer changes on their surface by stacking only two DNA base pairs into magnetic response that is reproducible and computable using the Fokker-Planck equation. By tuning the grafting density of DNA receptors on MARs, we identified two characteristic lengths: (1) the ssDNA receptor brush length of 22-24 nt in high density MARs and (2) the flanking domain length of > 13 nt that leads to unprecedented MARs properties. By varying the flanking domain length, we proposed the concept of coupled-decoupled binding-to-dynamics for the first time, where the flanking ssDNA segment decouples the effect of target binding from the magnetization dynamics. Using these concepts and exploiting DNA hairpins, we developed one-pot and one-design-for-all full combinatorial assays for sensing of four targets and the full combinatorial sets. By grafting dsDNA of varying lengths on MARs, we demonstrate that MARs are able to determine the average end-to-end distance of dsDNA with 1.3 ± 0.2 nm resolution up to ≈ 13 nm dynamic range and end-to-MAR distance of mono-streptavidin as a model protein system.

Exciting MARs with magnetic fields and reading out their signal magnetically open completely unexplored avenues for using molecular nanorulers in biology, as magnetic fields are compatible with biological entities and offer deep tissue penetration and less attenuated signal acquisition. Thanks to these unique features, we believe that MARs, by offering one-pot and rapid multiplexed sensing, resolving conformations of nucleic acid motifs in confined geometries, and determining protein-to-MAR molecular distances, will enable resolving molecular conformations of nucleic acids and proteins in complex biological environments such as extracellular matrix and intracellular in the near future.

## Associated Content

### Supporting Information

Experimental details including materials and methods (MARs synthesis and purification procedures, all the assay protocols, sample preparation and measurement methods), data analysis of fluorescence measurements, MPS, DLS and UV absorption measurements, all the fitting functions to analyze DLS and MPS data, Brownian Fokker-Planck simulation model description, AFM images, fluorescence intensity images and FCS analysis, Nupack simulation of DNA receptors used in this work, combinatorial biosensing of another set, and tables of all the DNA sequences applied in this work.

### Author contributions

A. L conceived the research idea. Y. W designed the research, performed surface modification, DNA labeling, assays, performed DLS and magnetic measurements, analyzed the data, prepared the graphics, and wrote the first draft of the manuscript. F. W developed and built the MPS setup. T. S performed single molecule fluorescence experiments and analyzed data. R. A synthesized nanoparticles. C. K conducted AFM imaging and analyzed the data. M. S, P. T, and J. L provided resources and analyzed data. T. V developed and built the custom MPS setup and performed the Fokker-Planck simulations. A. L provided resources, performed TEM measurements, supervised the research, analyzed data, and wrote the manuscript.

### Funding sources

Financial supports by the German Research Foundation (DFG, project number 545038923, A. L), Grant Numbers INST 86/2224-1 FUGG (project number 519922049, P. T & T. S), Metrology4life (A. L), the excellence clusters under Germany’s Excellence Strategy e-conversion-EXC 2089/1-390776260 and BioSysteM EXC3092/1-533751719, and the European Research Council Consolidator Grant “ProForce” (101002656) are acknowledged.

### Conflicts of interest

The authors declare no competing financial interest.

## Acknowledgments

We acknowledge Miss Kerstin Franke for laboratory support, Miss Petra Schmidt (TU Braunschweig) for ICP-OES measurements, Prof. Tim Liedl (LMU Munich) for giving the access to TEM measurements, and Dave J. van den Heuvel and Elleke van Harten for laboratory support.

## Notes

### Competing Interest Statement

The authors have declared no competing interest.

### Summary of Updates

The author list that appears on the website does not contain all the names.

