## Supporting Information for "Magnetic molecular nanorulers enable combinatorial biosensing and determine distances at the molecular level"

### Materials and Methods

#### Chemicals

Iron (III) acetylacetonate (99.9%, trace metal basis), cobalt (II) acetylacetonate (≥ 99.0%), dibenzyl ether (DBE, ≥ 98%), oleic acid (OA, 90%), 1-octadecene (ODE, 90%), poly(maleic anhydride-alt-1-octadecene) (Mn=30-50 kDa), $NH_2$-PEG(3)-Azide, Tris-Base, EDTA, NaCl, PBS buffer, and acetic acid were purchased from Sigma-Aldrich (Germany). Zinc (II) acetylacetonate (95%) was purchased from Merck. All ssDNA oligonucleotides were obtained from Biomers GmbH (Germany) with HPLC purification. Monovalent streptavidin was purchased from Merck. Sodium oleate (> 97%) was purchased from TCI (USA). All organic solvents with ACS reagent-grading were purchased from Carl Roth (Germany) and used without further purification.

#### Magnetic nanoparticle synthesis

Briefly, cobalt- and zinc-doped iron oxide nanocubes were synthesized via thermal decomposition of iron (III) acetylacetonate, cobalt (II) acetylacetonate, and zinc (II) acetylacetonate in a mixture of dibenzyl ether, 1-octadecene, oleic acid, and sodium oleate at 290°C for 30 min as previously described[1] with slight modifications[2].

#### Functionalization of poly(maleic anhydride-alt-1-octadecene) with $NH_2$-PEG(3)-azide linker

Poly(maleic anhydride-alt-1-octadecene) (PMAO) was modified with $NH_2$-PEG(3)-azide via anhydride ring opening reaction following the protocol published by Jin et al.[3] with moderate modifications. In a typical reaction, 338 mg (0.964 mmol monomer units) of PMAO were added into 17 mL of dimethylformamide (DMF) in a 50 mL round flask and heated up to 65°C while being stirred magnetically. Next, 191.3 µL (0.964 mmol) of NH2-PEG(3)-azide dissolved in 3 mL of DMF was pipetted into the reaction flask after the PMAO was completely dissolved. The reaction flask was then left to react for 24 h at 65°C. After cooling the flask to room temperature, the light yellowish and oily crude product was dialyzed against dichloromethane (DCM) using regenerated cellulose dialysis tubes (6 kDa cutoff size) to remove unreacted ingredients and byproducts. Finally, DCM was thoroughly removed in a rotary evaporator and the resulting polymers were dissolved in a specific amount of chloroform to reach a typical concentration of 35 mg/mL and stored at 4°C prior to usage.

**Polymer coating procedure**

The polymer coating of custom MNPs was performed using our and others protocols.[4–6] Typically, 2.64 mL (corresponding to 500 polymer units per $nm^2$ of a MNP) of PMAO-azide polymer (35 mg/mL) were diluted with 7.36 mL of chloroform in a 30 mL glass vial and sonicated for 10 min. Next, 2 mL of particle suspensions in chloroform (1.44 g(MNP)/L) were first diluted with 8 mL of chloroform and then added dropwise into the polymer solution while it was sonicated. The complete mixture was additionally sonicated for 10 min for homogenization. Further mixing was achieved by overnight shaking of the whole reaction mixture at room temperature. Next day, chloroform was removed slowly within 5-6 h through stepwise pressure reduction to a final value of 350 mbar and temperature increase to 34°C. After a complete removal of chloroform, particles were resuspended in 20 mL of sodium borate buffer (pH 8.7) by sonication for 1 h at ≈ 45°C. The particle suspension was then concentrated to 2 mL using spin filtration (Amicon regenerated cellulose spin filter, 15 mL, 50 kDa cut-off size) at 3500 rpm and 20°C for 20 min.

**Purification and particle fractionation**

To remove the excess of polymer, polymer micelles, and clusters that are formed during the polymer coating process, the polymer-coated MNPs were fractioned using non-continuous sucrose gradient (10%: 40%: 60%, from top to bottom, each fraction 4 mL) centrifuge columns, following our protocol.[6] Afterwards, the sucrose was removed through 3 rounds of Amicon filtration (Amicon spin filter, 15 mL, 50 kDa cutoff size) at 3500 rpm and 20°C for 13 min. After each round of centrifugation, the particles were thoroughly resuspended in borate buffer by vigorous pipetting and shaking. Finally, the MNPs were concentrated to 2 mL in borate buffer. To further remove the residual polymeric micelles, we performed one round of magnetic washing using a 1.5 mL MACS column (Miltenyi Biotec) in combination with a MiniMACS separator (permanent magnet) as reported earlier.[6] The MNPs were finally collected in 1.6 mL TE buffer (5 mM Tris-base, 5 mM NaCl, 1 mM EDTA, pH 7.3) and stored at room temperature for further processes.

**Agarose gel electrophoresis to fractionate polymer coated MNPs and obtain MARs**

To extract the fraction of MNPs that are encapsulated singly (monomeric MNPs) within a polymeric shell, native agarose gel electrophoresis (AGE) was applied. A 1% agarose gel was prepared by dissolving 0.5 g of agarose in 50 mL of 1 × TAE buffer by applying microwave irradiation for ≈ 2 min. After 5 min cooling, the homogenous gel solution was poured into a gel mold and allowed to solidify for 30 min. Afterwards, the casted gel was placed in electrophoresis chamber (Biorad) and samples were pipetted (containing 20 wt% sugar) into the wells. Next, AGE was performed in 1 × TAE running buffer at 90 mA for 30 min to extract the monomeric MNPs. The most advanced band (3 − 4 mm wide), corresponding to the monomeric MNPs, was subsequently excised from the gel. Finally, the particles were recovered by squeezing the gel fragments using parafilm coated glass slide. The monomeric polymer-coated MNPs are called magnetic molecular nanorulers (MARs).

**DNA functionalization of MARs**

The click-chemistry conjugation enables grafting ssDNA onto MARs. In a typical labeling, a 150 µL of MARs suspension in TE buffer at a particle concentration of 15 nM was mixed with 32.8 µL, 16.4 µL, 8.2 µL or 4.1 µL of 100 µM oligos of interest in TE buffer to obtain 1.2, 0.6, 0.3 or 0.1 ssDNA/nm$^2$ nominal grafting density. After complete mixing, the sample was incubated at room temperature overnight. To maximize the labeling efficiency of ssDNA to MARs, we applied salt aging procedure using 5 M NaCl the next day by increasing the NaCl concentration in the labeling reaction incrementally by 100 mM every hour and up to 400 mM. After each salt adjustment step, the mixture was homogenized by vortexing and sonicating for 20 s. At the end of the salt aging procedure, the mixture was again incubated at room temperature overnight. Next day, excess of ssDNA was washed out by 2 rounds of centrifugation at 13.000 × g and 5°C for 12 min. Finally, the sample volume was adjusted to 150 µl to obtain the original particle concentration. The samples were stored at 4°C prior to further use.

Prior to labeling MARs with loop-receptor DNA strands for combinatorial assays, an annealing process was conducted by incubating ssDNA receptors (100 µM) in TE buffer (pH = 7.3) at 95°C for 5 min and then cooling to 25°C at 2°C per min. Subsequently, DNA functionalization was performed as described above.

**Hybridization assays: target DNA to ssDNA-labelled MARs**

For the titration assays, MARs functionalized with ssDNA receptors were first diluted to 1.5 nM and mixed with complementary target DNA (tDNA) at 150 mM NaCl in TE buffer (pH 7.3). The titration was performed by incrementally increasing the molar ratio of tDNA to receptor ssDNA ($R_{t/r}$) as follows: 0.05:1, 0.1:1, 0.2:1, 0.4:1, 0.8:1, 1:1 and 2:1. The reaction mixtures was then incubated for 24 h in a thermomixer at 25°C with 10 min shaking intervals (5 min shaking at 550 rpm and 5 min no shaking). For all the following assays, the tDNA was added at the ratio of $R_{t/r}$ = 2:1 using the same buffer composition and reaction conditions described above. The excess ratio ensures that all available ssDNA receptors are hybridized to target strands.

**Binding monovalent streptavidin to biotinylated dsDNA-labelled MARs**

To mitigate MARs crosslinking arising from the four tetravalent binding sites of conventional streptavidin, we utilized monovalent streptavidin (monostreptavidin) and 20% biotinylated dsDNA labelled on MARs. After labeling MARs with various ssDNA receptors ranging from 9 to 80 nt (Table S3) and washing unbound ssDNA, we added target DNA (20% biotinylated: 80% non-biotinylated) at the ratio of $R_{t/r}$ = 2:1 (target/receptor). The reaction mixture was next incubated for 24 h in a thermomixer at 25°C with 10 min shaking intervals (5 min shaking at 550 rpm and 5 min no shaking). Afterwards, the excess target DNA was removed by one round of centrifugation at 13.000 × g and 5°C for 10 min. The dsDNA-labelled MARs were next suspended in PBS buffer (pH = 7.4) for monostreptavidin binding. To saturate all available biotin molecules on MARs, streptavidin in PBS was added to MARs at a 5:1 molar ratio relative to biotin. The reaction mixtures were then incubated overnight in a thermomixer at 25°C with 10 min shaking intervals (5 min shaking at 550 rpm and 5 min no shaking).

**Sample purification for fluorescence lifetime measurements on MARs**

To remove unbound ssDNA-dye strands that would otherwise interfere with the signal from the bound dyes, the MARs were purified through centrifugation at 6.000 × g for 1 minute using Amicon spin filter (0.5 mL, 100 kDa cutoff size). A 400 μL of TE buffer (pH = 7.3) was added into the purified sample and the mixture was homogenized by gentle pipetting within the filter chamber. This washing procedure was repeated seven times to completely remove unbound dyes. Subsequently, the filter was flipped upside down into a new clean tube and centrifuged for 1 minute at 2.000 × g to recover the MARs. The samples were stored at 4°C until further use.

**Transmission Electron Microscopy (TEM)**

Transmission electron microscopy measurements were carried out using a JEOL TEM microscope operating at 100 kV. The TEM samples were prepared by drop-casting 5 μl of particle chloroform suspensions on a grid (Formvar-carbon coated copper grids, mesh size of 300) and letting the chloroform completely evaporate in a fume hood. Negative-stain TEM samples were prepared by staining with uranyl formate (1%) for 10 s after incubating particle water suspensions on a grid for 5 min. The particle core size $D_c$ was determined by analyzing large fields of view of TEM images using the automatic particle analysis function of the ImageJ software.

**Dynamic Light Scattering (DLS)**

DLS measurements were performed using a Malvern Zetasizer instrument at 173° backscattered measurement mode. The measured samples for DLS are typically 60 μL of particle aqueous suspensions at a particle concentration of 0.015-0.02 g/L at 25°C.

**Atomic Force Microscopy (AFM) imaging**

The measurements were performed on a Nanowizard Ultraspeed 2 high speed AFM (JPK) with triangular FASTSCAN A probes (Bruker) with a driving frequency of ~ 1400 kHz and a spring constant of 18 N/m. The samples were scanned in the tapping mode in air over different fields of view of 5 × 5 $\mu m^2$ with a resolution of ~ 1 nm/pixel with a line scanning speed of 1 Hz. For AFM imaging, 20 μL of particles in TE buffer containing 300 mM NaCl were deposited onto a freshly cleaved mica substrate and incubated for 15 min in a sealed chamber, followed by rinsing with 20 mL of Milli-Q water and drying under a gentle stream of pure nitrogen gas. The obtained AFM images were processed in MountainsSPIP 10 analysis software (Digital Surf) using line-by-line correction and polynomial background subtraction. The height of individual nanoparticles was extracted with the particle and pore analysis tool of the software.

**Magnetic Particle Spectroscopy (MPS)**

MPS measurements were carried out on a custom-built setup, which is designed for sensitive magnetic bioassays and tailored to MARs magnetic and hydrodynamic characteristics.[7] The MPS operates at an excitation frequency of 1954 Hz and a magnetic field amplitude of $\mu_0 H$ = 15 mT. The measurements were performed on 60 μl particle aqueous suspensions at a typical particle concentration of 1.5 nM and at 22-25°C.

**Solution based single particle fluorescence microscopy for fluorescence lifetime measurements**

The fluorescence lifetime measurements on MARs were performed on the commercial confocal microscope Luminosa (PicoQuant GmbH). For solution experiments, the organic dyes were excited by a pulsed laser at 640 nm wavelength, operating at 40 MHz with a power of 5 μW. The laser light was guided into the epi-illuminated confocal microscope by dual-band beam splitter

(DB485/640) and focused by an oil immersion objective (100×, NA 1.45, Olympus) into the sample. The emitted fluorescence was collected through the same objective and spatially filtered by a pinhole with a 50 µm diameter. The fluorescence signal was cleaned by bandpass filters (BP690/70) and focused on single photon counting module. A second APD (avalanche photodiode) was also recorded with a bandpass (BP582/64) filter in the detection path for the software analysis which requires a second detector, even though no meaningful signal but uncorrelated signal is detected. The measurements were conducted in a home-built 60 µL chamber on 170 µm thick 1.5H glass slide. The chamber and glass surface were passivated with a 1 mg/mL BSA solution for 2 min to prevent unspecific particle sticking to surface. Afterwards, the chamber was rinsed three times with the storing buffer. The MARs were added to the buffer on the microscope, and the laser was focused 5 µm deep into the solution. The concentration of MARs was adjusted on the microscope, so that the bursts were well separated in the fluorescence transient of 1 ms binning. Subsequently, multiple fluorescence transients were recorded with a length of 10 min at the same spot for a total acquisition time of 130 min.

**Data analysis on fluorescence lifetime measurements**

Data analysis was performed with the open source software Pulsed interleaved excitation Analysis with MATLAB (PAM).[8] Single-particle events were identified using a two-channel APBS-algorithm with a threshold of 5 photons per 500 µs time window and a minimum number of 30 photons per burst. For each burst, the lifetime was fitted with the implemented reconvolution fit. Bursts were filtered in the Burst Browser. The plotting of fluorescence lifetime vs. photons per burst reveal two lifetime populations, where the long fluorescence lifetime refers to the unquenched fluorescence lifetime of the ATTO643 dye and the quenched component to the dyes on MARs. As higher photon numbers per burst result in less uncertainty of the fluorescence lifetime, only bursts with more than 300 photons were used for building the fluorescence lifetime decays in figure 1f and S3b. To calculate the hydrodynamic size $D_h$ of MARs based on the diffusion coefficient $D$, the Stokes-Einstein-Sutherland equation[9] was applied:

$$D = \frac{k_B T}{6\pi\eta R_H} \qquad \text{Eq.1}$$

Here, $k_B$ is the Boltzmann constant, $T$ denotes the absolute temperature, $\eta$ represents the fluid shear viscosity (taken as the dynamic viscosity of water at 25°C, $0.89\times10^{-3}$ kg/(m·s)), and $R_h$ is the hydrodynamic radius of the sphere particles ($R_h = D_h/2$).

**Fluorescence lifetime imaging microscopy of immobilized MARs**

Surface immobilization experiments with MARs were performed in Lab-Tek chambers using BSA–biotin–NeutrAvidin chemistry. Prior to use, the chambers were cleaned with 200 µL of Hellmanex (Hellma GmbH & Co. KG) for 2 h, rinsed with deionized water, and subsequently washed with 200 µL of 1× PBS. The chambers were then incubated with 100 µL of biotinylated BSA (1 mg/mL) for 1 min, followed by a single washing step with 200 µL of 1× PBS. Subsequently, 100 µL of NeutrAvidin (1 mg/mL) was added and incubated for 1 min, followed by another washing step with 200 µL of 1× PBS. To enable MAR immobilization, 100 µL of a biotinylated DNA oligonucleotide (1 µM) with a sequence reverse-complementary to the DNA strand of the MARs was added and incubated for 2 min. The chambers were then washed five times with 200 µL of 1× PBS.

To facilitate hybridization of the MARs to the surface-immobilized ssDNA, Tris buffer containing 500 mM NaCl was added to screen the electrostatic repulsion between the DNA strands. The purified MAR sample was subsequently introduced into the chamber, and the surface density was monitored using a Luminosa microscope. After 10 min, a sufficient surface density was achieved, and the chambers were washed three times with TE buffer to remove unbound MARs. For subsequent fluorescence lifetime imaging microscopy (FLIM) measurements, the chambers were filled with TE buffer.

**MPS-based magnetic bioassays and analysis of MPS harmonic ratio**

Our magnetic assays measure changes in the magnetic Brownian relaxation time of MNPs in free solution, a characteristic relaxation mechanism describing particles physical rotation. When the particle hydrodynamic size/volume increases upon a molecular recognition between surface receptors and target molecules, e.g. nucleic acids and proteins, the rotation slows down and the relaxation time increases. These changes are reflected in the MPS harmonics spectrum. A molecular binding slows down the Brownian relaxation of MNPs in ac magnetic fields and thereby reduces the MPS harmonics amplitudes. The MPS higher harmonics are intrinsically more sensitive to changes in the particle relaxation dynamics than the fundamental excitation frequency, which means that the $5^{th}$ harmonics drops more than the $3^{rd}$ harmonics in response to binding. Therefore, the ratio of the $5^{th}$ to $3^{rd}$ harmonics (HR53) drops further after binding/sensing of the target molecule compared to that of the sample without the target molecule. Although the individual harmonics amplitudes depend on the particle magnetic mass, their ratio is a particle concentration-independent metric to judge the success of MPS-based bioassays.

**Determination of DNA grafting density on MARs**

To determine the ssDNA grafting density on MARs, UV absorption measurement at 260 nm was performed on supernatants (from each round of centrifugation) that were obtained from DNA labeling samples after pelleting the MARs. Following our procedure,[6] we quantified the number of grafted ssDNA per MAR ($D_h$ = 28 nm) and computed its corresponding grafting density (Table S1). The grafting densities $\rho_{DNA}$ are calculated using:

$$\rho_{\text{DNA}} = \frac{\text{ssDNA per MAR}}{S_{\text{MAR}}} \qquad \text{Eq. 2}$$

**Hill equation for fitting $D_h$ vs. grafting density and titration assays**

Volume-weighted hydrodynamic size $D_h$ was fitted with a Hill-like equation given by:

$$D_h(x) = A_1 + (A_2 - A_1)\frac{x^n}{(x^n + k_d{}^n)} \qquad \text{Eq. 3}$$

$A_1$, $A_2$, $k_d$, and $n$ are fitting parameters. $k_d$ indicates the DNA grafting density that generates a half-maximum increase in $D_h$ and $n$ is the Hill coefficient reflecting the cooperativity.

To quantify the titration curves (Fig. 2) at each DNA grafting density, a similar Hill equation was applied given by:

$$HR53_{(x)} = HR53_{(start)} + \left(HR53_{(end)} - HR53_{(start)}\right)\frac{c_t{}^n}{c_t{}^n + k_d{}^n} \qquad \text{Eq.4}$$

$k_d$, and $n$ are fitting parameters, which represent the apparent affinity of target to receptor DNA on MARs and cooperativity of the system, respectively. $c_t$ is the target DNA concentration.

**Relative HR53 changes and their normalization to the grafted DNA number**

To better compare the effects of DNA grafting density on the magnetic response of MARs, we calculated the HR53 relative changes ΔHR53 in % determined via:

$$\Delta HR53\ (\%) = 100 \times \frac{HR53_{ctrl} - HR53_t}{HR53_{ctrl}} = 100 \times (1 - \frac{HR53_t}{HR53_{ctrl}}) \qquad \text{Eq. 5}$$

$HR53_{ctrl}$ and $HR53_t$ are the harmonic ratios of control sample without complementary DNA and the titration sample with $R_{t/r}$ = 2:1, respectively. ΔHR53 (%) is normalized to the number of grafted DNA strands at respective densities (Table S1).

**Brownian Fokker-Planck simulation model**

The influence of dsDNA shell thickness on the dynamic magnetic response was modeled using a rotational Brownian Fokker–Planck (FP) approach.[10–12] The magnetic moment was assumed to be fixed to the particle's magnetization axis, such that magnetic relaxation occurs by physical rotation of the entire particle (i.e. Brownian mechanism) in solution. The orientational probability density P(u,t) evolves according to

$$2\tau_B \frac{\partial P}{\partial t} = \nabla_{\mathbf{u}} \cdot [\nabla_{\mathbf{u}} P + P \nabla_{\mathbf{u}} U(\mathbf{u}, t)], \qquad \text{Eq.6}$$

where $\mathbf{u}$ is the particle orientation, $\nabla_{\mathbf{u}}$ denotes the surface gradient on the unit sphere, and $U(\mathbf{u}, t)$ is the magnetic energy normalized by $k_B$T. For an applied field $\mathbf{H}(t)$,

$$U(\mathbf{u}, t) = -\frac{\mu_0 m \mathbf{H}(t) \cdot \mathbf{u}}{k_B T}, \qquad m = M_s V_c. \qquad \text{Eq.7}$$

The Brownian relaxation time is determined by the hydrodynamic particle volume,

$$\tau_B = \frac{3\eta V_h}{k_B T} = \frac{\eta \pi D_h^3}{2 k_B T}. \qquad \text{Eq. 8}$$

Shell-thickness variations were introduced through the hydrodynamic diameter,

$$D_h = D_{h,0} + 2R_{\text{em}}, \qquad \text{Eq. 9}$$

where $D_{h,0}$ represents the hydrodynamic diameter of MARs without dsDNA, which was determined to be 39.6 nm using DLS and $R_{em}$ represents the shell thickness increase, which is equivalent to the end-MAR distance of dsDNA on particles. The magnetic core volume and saturation magnetization were kept constant at 6434 $nm^3$ and 291 kA/m (determined from VSM magnetometry), respectively. Thus, shell growth affects the simulated response through the cubic dependence of $\tau_B$ on $D_h$, thereby changing the rotational dynamics without directly modifying the particle magnetic moment.

For sinusoidal excitation, the periodic FP solution was obtained by expanding the orientational distribution in Legendre modes and solving the resulting harmonic system using a truncated continued-fraction scheme. Magnetization harmonics and harmonic ratios were calculated from the periodic steady-state response; for polydisperse samples, the complex harmonic response was averaged over the particle distribution before calculating harmonic magnitudes and ratios. In this work, we assumed lognormal distribution for particle core size and hydrodynamic size of MARs with the standard deviation σ of 0.1 and 0.15, respectively.

The model is applicable to Brownian relaxing MNPs, as it is the case for our cobalt containing particles, and to particles that can rotate freely in solution. Particle immobilization, viscoelastic matrix torques, interparticle interactions, and coupled Brownian-Néel relaxation are not included in our modelling.

**Determination of the DNA shell thickness based on Z-average values obtained from DLS**

We verified that the experimental HR53 values are highly consistent with the simulated values from the PF simulations when DLS Z-average sizes are taken as particle hydrodynamic sizes. Therefore, all the calculations regarding shell thickness are based on Z-average values.

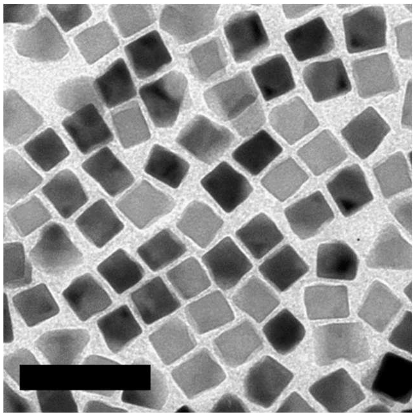

**Figure S1.** Negative-stain transmission electron microscopy (TEM) image of our custom $Co_{0.4}Zn_{0.2}Fe_{2.4}O_4$ MNPs in chloroform. The scale bar is 60 nm.

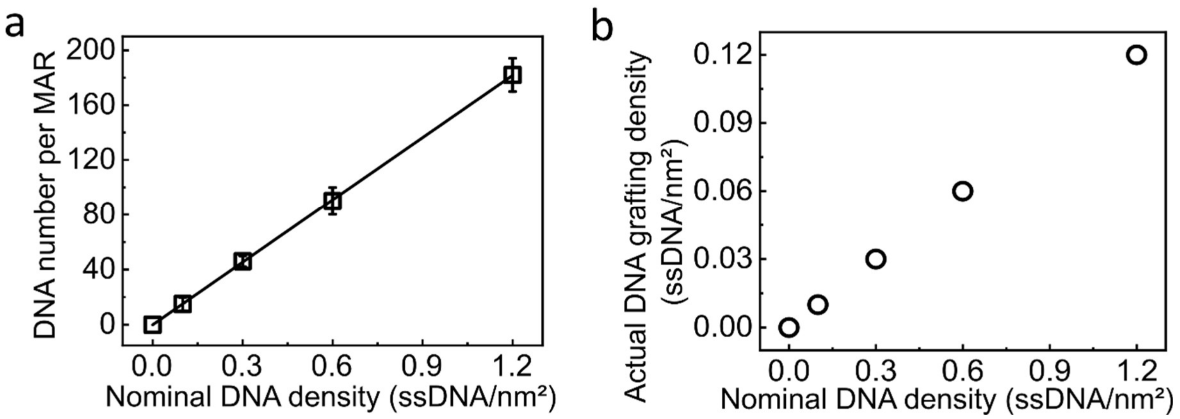


**Figure S2.** (a) The average ssDNA number per MAR ± standard error of the mean (SEM) ($n = 3$) as estimated from the depletion assay vs. the nominal DNA grafting density. (b) The actual DNA grafting density as calculated based on DNA number per MAR vs. nominal grafting density (see also in Table S1). Unless otherwise noted, actual DNA densities are plotted in all main-text figures.

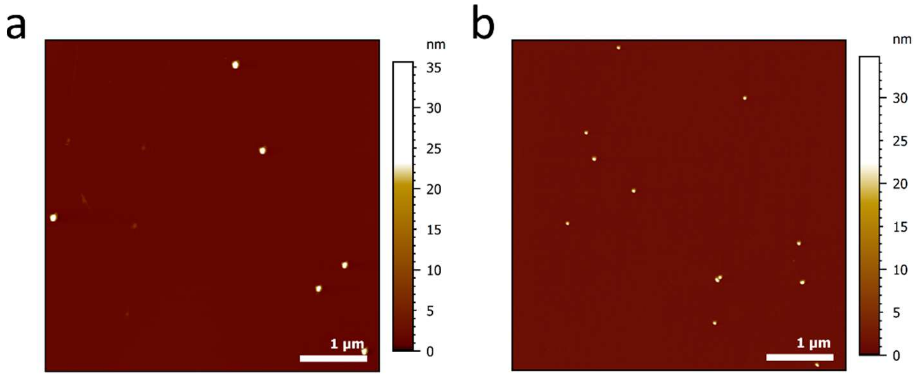


**Figure S3.** Representative atomic force microscopy (AFM) images of MARs at (a) 0.03 and (b) 0.06 ssDNA/nm$^2$ density. The images show single nanoparticles with a height distribution quantified in Figure 1f.

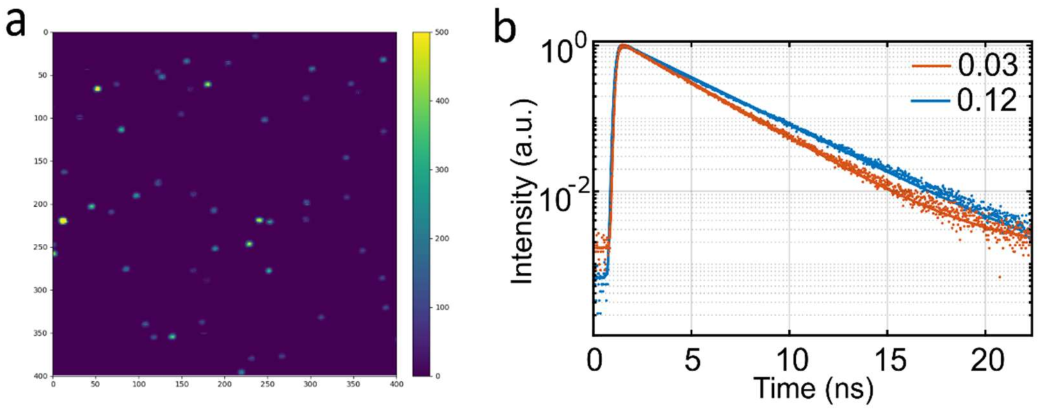


**Figure S4.** (a) The fluorescence intensity image of Fig. 1h at 0.12 ssDNA/nm² density of DNA receptor 39-1 sparsely labeled with ATTO643. An upper intensity threshold of 500 photons per pixel was applied. The scale bar on the left and bottom axis depict the number of pixels where each pixel has a length of 50 nm. (b) Fluorescence lifetime decays of the quenched ATTO643 dye labeled MARs populations at 0.03 and 0.12 ssDNA/nm², which are cumulative data of Fig. 1g. Only bursts with a fluorescence lifetime in the interval from 2.0 to 3.4 ns were considered.

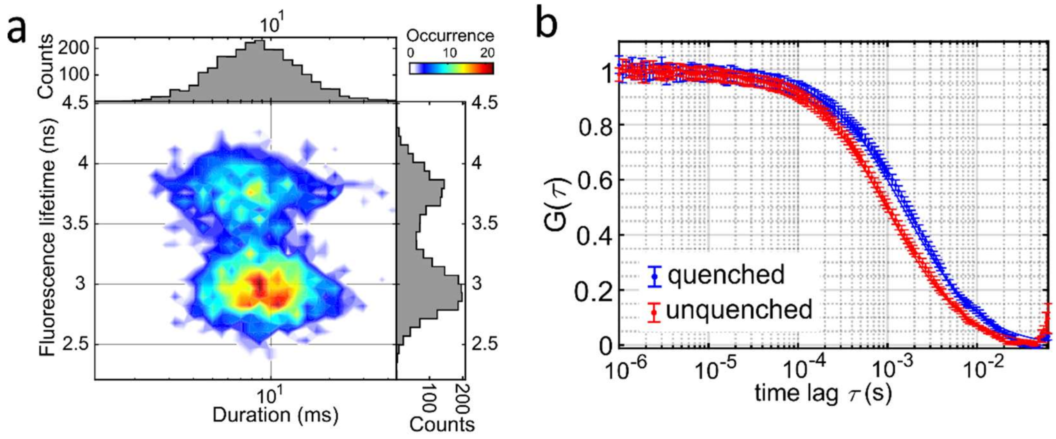


**Figure S5.** (a) The two-dimensional correlation plot of fluorescence lifetime vs. dwell time on MARs at 0.12 ssDNA/nm² density in Figure 1g. The slight shift in dwell time indicates distinct diffusion coefficients and therefore hydrodynamic radii of two populations. The color scale (top right) represents event occurrence, which corresponds to the counts plotted on the right and top. (b) Fluorescence correlation spectroscopy (FCS) analysis of (a). Diffusion coefficients of 8.75 μm²/s and 13.53 μm²/s were extracted from the quenched (blue) and unquenched (red) populations, respectively.

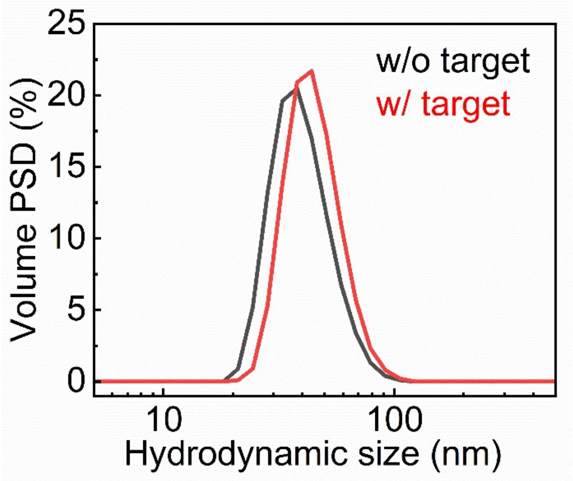


**Figure S6.** Volume-weighted particle size distributions of MARs at 0.12 density before and after adding fully complementary target DNA as measured by dynamic light scattering (DLS).

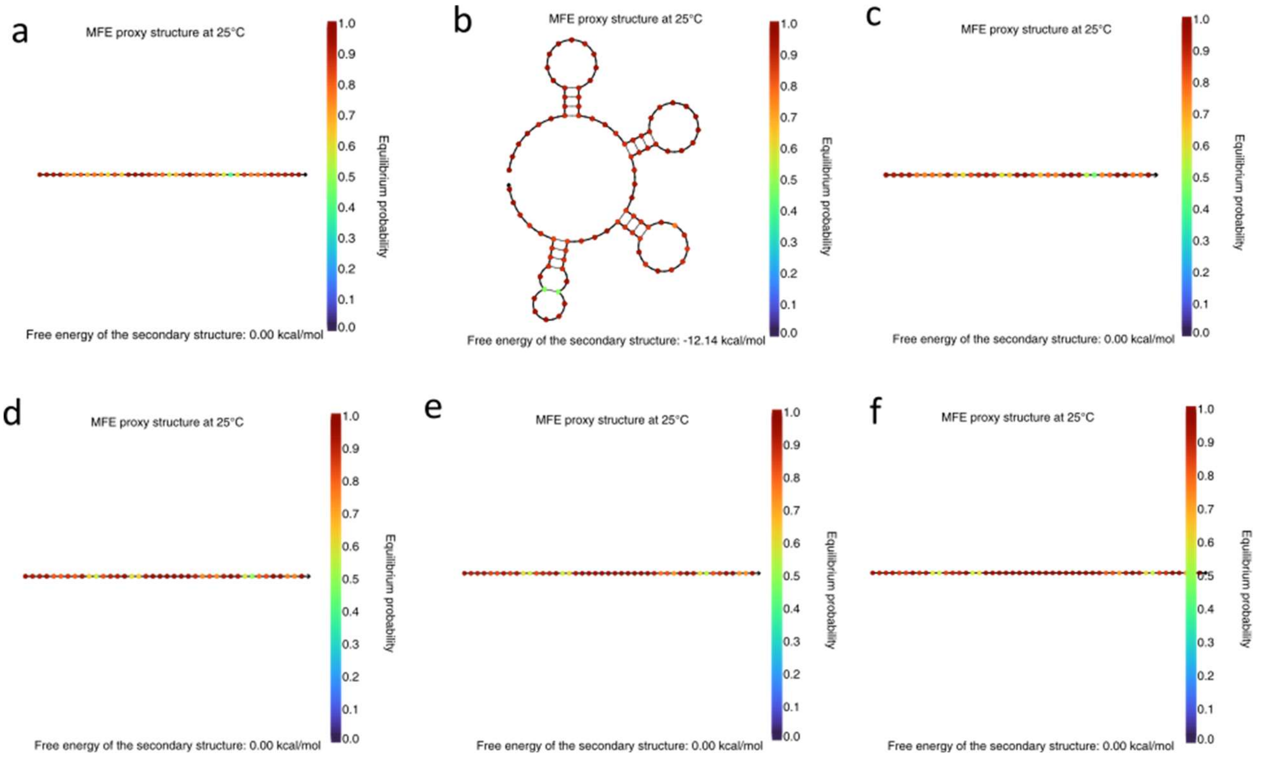


**Figure S7.** NUPACK simulations of potential secondary structures of (a) receptor 39-2, (b) loop-receptor, (c) 2D-0T-receptor, (d) 2D-5T-receptor, (e) 2D-10T-receptor, and (f) 2D-15T-recptor.[13] None of the DNA receptor structures except for the loop receptor exhibit secondary structures.

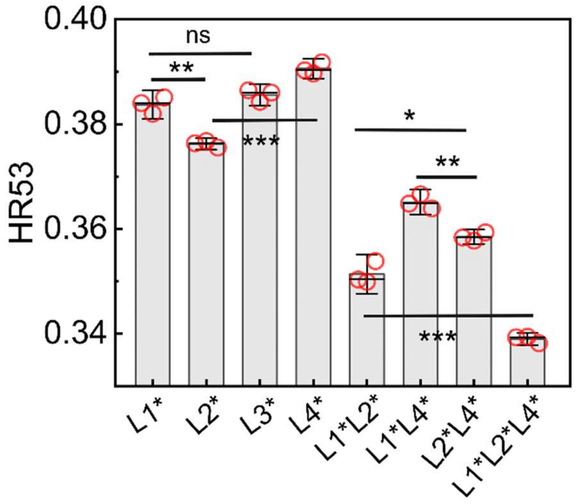


**Figure S8.** Zoomed-in view of the full combinatorial cases of L1*, L2* and L4* plus single L3* target binding response from Fig. 5c set 1, demonstrating that three targets L1*, L2*, L4* and their random combinations are with high significance distinguishable. The p values were determined using unpaired two-tailed t-tests (*$p < 0.05$, **$p < 0.01$, ***$p < 0.001$; ns, not significant). The grafting density applied was 0.12. Error bars represent 3 × SEM of three independent biological replicates. The measurements were performed on samples with a typical particle concentration of 1.5 nM.

**Table S1. Overview of nominal and actual grafting densities**

| Nominal DNA grafting density / ($nm^2$ of MAR) | ssDNA per MAR | Actual DNA grafting density / ($nm^2$ of MAR) |
|---|---|---|
| 0.1 | 16 ± 2.4 | 0.01 |
| 0.3 | 46 ± 3.5 | 0.03 |
| 0.6 | 90 ± 5.2 | 0.06 |
| 1.2 | 182 ± 8.8 | 0.12 |

**Table S2.** DNA oligonucleotides used in the titration assays and spatial bindings at the proximal end and the distal end of DNA receptors.

| Name | Sequence (5'→ 3') | Length (nt) |
|---|---|---|
| Receptor 39-1 | 5'DBCO-AAA AAA GGA TTG CGG GTG CCA ATG TGA TCT TTT GGT GTA | 39 |
| Receptor 39-2 | 5'DBCO-AAT TAA GAG ATA CGG TAG ACA ATG TGA TCT TTT GGT GTA | 39 |
| Target 39-1 | TAC ACC AAA AGA TCA CAT TGG CAC CCG CAA TCC T | 34 |
| Target-P-12 | TAC CGT ATC TCT | 12 |
| Target-P-13 | CTA CCG TAT CTC T | 13 |
| Target-P-14 | TCT ACC GTA TCT CT | 14 |
| Target-P-16 | TGT CTA CCG TAT CTC T | 16 |
| Target-P-18 | ATT GTC TAC CGT ATC TCT | 18 |
| Target-P-20 | ACA TTG TCT ACC GTA TCT CT | 20 |
| Target-P-22 | TCA CAT TGT CTA CCG TAT CTC T | 22 |
| Target-P-24 | GAT CAC ATT GTC TAC CGT ATC TCT | 24 |
| Target-P-26 | AAG ATC ACA TTG TCT ACC GTA TCT CT | 26 |
| Target-P-28 | AAA AGA TCA CAT TGT CTA CCG TAT CTC T | 28 |
| Target-P-30 | CCA AAA GAT CAC ATT GTC TAC CGT ATC TCT | 30 |
| Target-P-32 | CAC CAA AAG ATC ACA TTG TCT ACC GTA TCT CT | 32 |
| Target-39-2 | TAC ACC AAA AGA TCA CAT TGT CTA CCG TAT CTC T | 34 |
| Target-D-12 | TAC ACC AAA AGA | 12 |
| Target-D-13 | TAC ACC AAA AGA T | 13 |
| Target-D-14 | TAC ACC AAA AGA TC | 14 |
| Target-D-16 | TAC ACC AAA AGA TCA C | 16 |

| | | |
|---|---|---|
| Target-D-18 | TAC ACC AAA AGA TCA CAT | 18 |
| Target-D-20 | TAC ACC AAA AGA TCA CAT TG | 20 |
| Target-D-22 | TAC ACC AAA AGA TCA CAT TGT C | 22 |
| Target-D-24 | TAC ACC AAA AGA TCA CAT TGT CTA | 24 |
| Target-D-26 | TAC ACC AAA AGA TCA CAT TGT CTA CC | 26 |
| Target-D-28 | TAC ACC AAA AGA TCA CAT TGT CTA CCG T | 28 |
| Target-D-30 | TAC ACC AAA AGA TCA CAT TGT CTA CCG TAT | 30 |
| Target-D-32 | TAC ACC AAA AGA TCA CAT TGT CTA CCG TAT CT | 32 |

**Table S3.** DNA oligonucleotides designed for 2-plex assays and combinatorial biosensing.

| Name | Sequence (5’→ 3’) | Length (nt) |
|---|---|---|
| Loop-receptor | 5’DBCO-AAA CGC CCA CTC TTA GTC TGT GGA TCC CAG ATC TTC ACA CTG GTT AGG CTT ACG GAT TCA GCC ATA GGA CTT CTA CTA GGT CCG TAC G | 88 |
| Target-L1* | ATC CAC AGA CTA AGA GTG GG | 20 |
| Target-L2* | AAC CAG TGT GAA GAT CTG GG | 20 |
| Target-L3* | ATG GCT GAA TCC GTA AGC CT | 20 |
| Target-L4* | ACG GAC CTA GTA GAA GTC CT | 20 |
| 2D-receptor | 5’DBCO-AAT TAT CAT ACA TAC GTC TTG CAT TCT GAT TCC AT | 35 |
| 2D-5T-receptor | 5’DBCO-AAT TAT CAT ACA TAC GTC TTT TTT TGC ATT CTG ATT CCA T | 40 |
| 2D-10T-receptor | 5’DBCO-AAT TAT CAT ACA TAC GTC TTT TTT TTT TTT GCA TTC TGA TTC CAT | 45 |
| 2D-13T-receptor | 5’DBCO-AAT TAT CAT ACA TAC GTC TTT TTT TTT TTT TTT GCA TTC TGA TTC CAT | 48 |
| 2D-15T-receptor | 5’DBCO-AAT TAT CAT ACA TAC GTC TTT TTT TTT TTT TTT TTG CAT TCT GAT TCC AT | 50 |
| 2D-20T-receptor | 5’DBCO-AAT TAT CAT ACA TAC GTC TTT TTT TTT TTT TTT TTT TTT TGC ATT CTG ATT CCA T | 55 |
| 9bp-receptor | 5’DBCO-AAT TAT CAT ACA TA | 14 |
| 11bp-receptor | 5’DBCO-AAT TAT CAT ACA TAC G | 16 |
| 15bp-recptor | 5’DBCO-AAT TAT CAT ACA TAC GTC TT | 20 |
| 3D-receptor | 5’DBCO-AAT TAT CAT ACA TAC GTC TTG CAT TCT GAT | 50 |

| | | |
|---|---|---|
| | TCC ATA GTA TTA TAC CTA TT | |
| 60bp-receptor | 5’DBCO-AAT TAT CAT ACA TAC GTC TTG CAT TCT GAT TCC ATA GTA TTA TAC CTA TTA CAC TGA TAT TTA CT | 65 |
| 70bp-receptor | 5’DBCO-AAT TAT CAT ACA TAC GTC TTG CAT TCT GAT TCC ATA GTA TTA TAC CTA TTA CAC TGA TAT TTA CTC TTA CAG ACT | 75 |
| 9bp-target | (5’biotin-) TAT GTA TGA | 9 |
| 11bp-target | (5’biotin-) CGT ATG TAT GA | 11 |
| Target-D1* | (5’biotin-) AAG ACG TAT GTA TGA | 15 |
| Target-D2* | (5’biotin-) ATG GAA TCA GAA TGC | 15 |
| Target-D3* | (5’biotin-) AAT AGG TAT AAT ACT | 15 |
| Target-D4* | (5’biotin-) AGT AAA TAT CAG TGT | 15 |
| Target-D4L* | (5’biotin-) AGT CTG TAA GAG TAA ATA TCA GTG T | 25 |
